# High-Resolution Chemical Mapping and Microbial Identification of Rhizosphere using Correlative Microscopy

**DOI:** 10.1101/2021.02.05.429689

**Authors:** Chaturanga D. Bandara, Matthias Schmidt, Yalda Davoudpour, Hryhoriy Stryhanyuk, Hans H. Richnow, Niculina Musat

**Affiliations:** Department of Isotope Biogeochemistry, Helmholtz-Center for Environmental Research (UFZ), Permoserstraße15, 04318 Leipzig, Germany

**Keywords:** CARD-FISH, Chemical microscopy, Fluorescence microscopy, HIM, Nano-SIMS, Resin embedding, Rhizosphere, Soil Bacteria, Spatial distribution, Water-jet cutting

## Abstract

During the past decades, several stand-alone and combinatory methods have been developed to investigate the chemistry (i.e. mapping of elemental, isotopic and molecular composition) and the role of microbes in soil and rhizosphere. However, none of these approaches are currently capable of characterizing soil-root-microbe interactions simultaneously in their spatial arrangement. Here we present a novel approach that allows chemical and microbial identification of the rhizosphere at micro-to nano-meter spatial resolution. Our approach includes i) a resin embedding and sectioning method suitable for simultaneous correlative characterization of *Zea mays* rhizosphere, ii) an analytical work flow that allows up to six instruments/techniques to be used correlatively, and iii) data and image correlation. Hydrophilic, immunohistochemistry compatible, low viscosity LR white resin was used to embed the rhizosphere sample. We employed waterjet cutting and avoided polishing the surface to prevent smearing of the sample surface at nanoscale. Embedding quality was analyzed by Helium Ion Microscopy (HIM). Bacteria in the embedded soil was identified by Catalyzed Reporter Deposition-Fluorescence *In Situ* Hybridization (CARD-FISH) to avoid interferences from high levels of auto fluorescence emitted by soil particles and organic matter. Chemical mapping of the rhizosphere was done by Scanning Electron Microscopy (SEM) with Energy-dispersive X-ray analysis (SEM-EDX), Time-of-Flight Secondary Ion Mass Spectrometry (ToF-SIMS), nano-focused Secondary Ion mass Spectrometry (nanoSIMS), and confocal Raman spectroscopy (µ-Raman). High-resolution correlative characterization by six different techniques followed by image registration shows that this method can meet the demanding requirements of multiple characterization techniques to chemically map the rhizosphere and identify spatial organization of bacteria. Finally, we presented individual and correlative workflows for imaging and image registration to analyze data. We hope this method will be a platform to combine various 2D analytics for an ample understanding of the rhizosphere processes and their ecological significance.

**Graphical Abstract:** 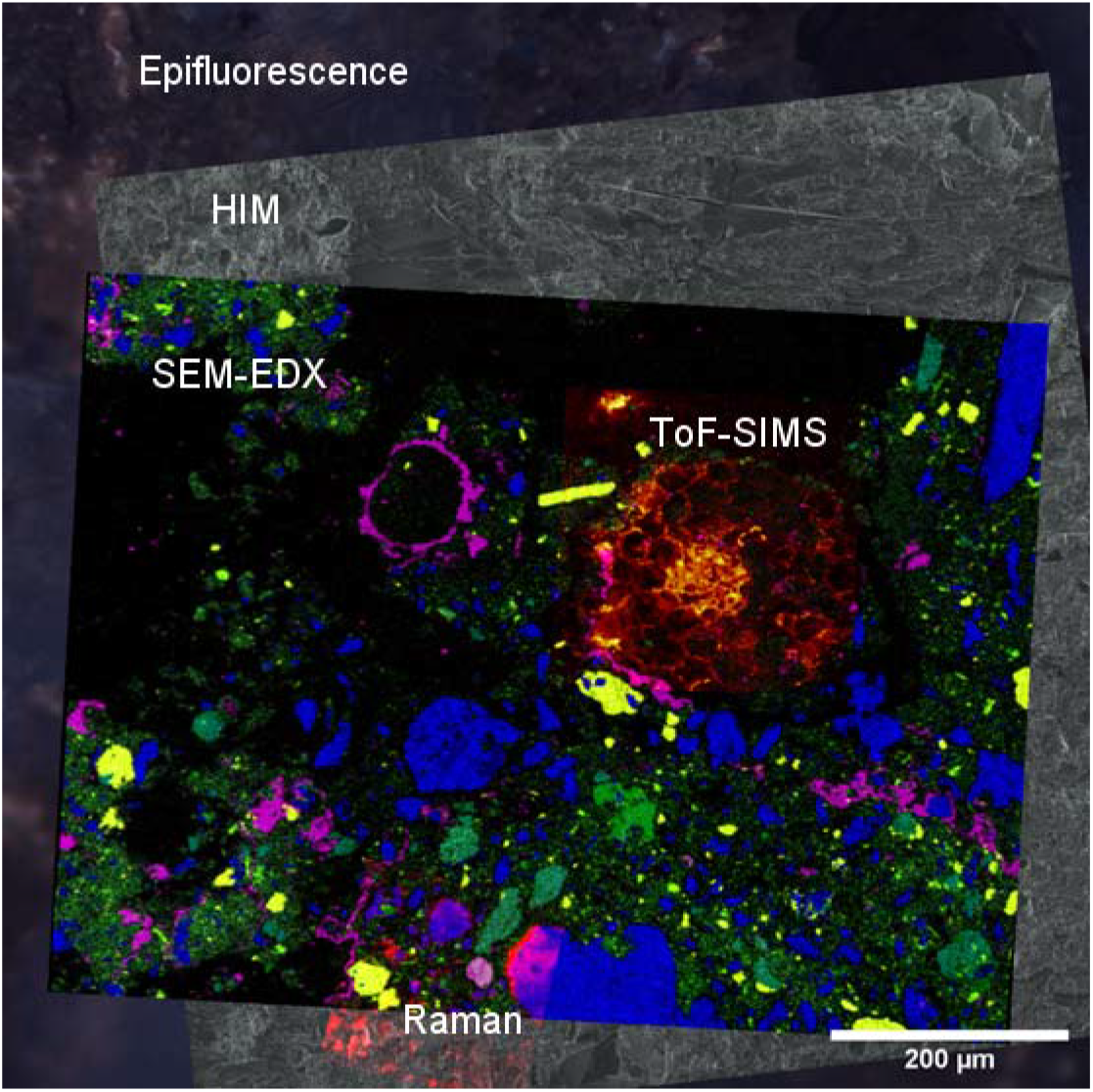

## Introduction

Soils are heterogeneous mixtures of various organic materials, mineral particles, and pores, which provides a matrix for plant growth and resource recycling by microbiome. Soil structure defines microenvironmental conditions and processes such as water retention, root penetration, microbial distribution, nutrient and gas exchange.^1-3^ Though these individual aspects are extensively studied, correlative studies relating each other are limited. Particularly, the structural and biochemical complexity of the rhizosphere as the main driver of terrestrial biogeochemical cycles is little understood. To explore such relationships between multiple components,^4^ it is crucial to characterize all soil components keeping its structural and chemical integrity i.e. chemical composition and spatial organization of minerals, non-living organic matter and microbial counterparts such as bacteria, fungi, and plant roots during the analysis.^5-12^ Main reasons that confine studies are associated with sample preparation to accommodate for multiple techniques allowing the concomitant analysis of; 1) root 2) soil and 3) microbes in their spatial context. A comprehensive chemical characterization coupled to microbial identification and distribution at spatial scales relevant for biogeochemical processes are necessary to achieve a systematic understanding of the key factors governing the self-organization and resilience.^11, 12^

Of particular interest in the rhizosphere research are the inherent microorganisms, key players of the carbon and nutrient recycling processes,^13, 14^ for their correct identification and quantification, fluorescent dyes should stain all microbial cells independent of their physiological state and metabolic activity.^15, 16^ In an environmental sample, unspecific binding sites for fluorescence stains are significantly higher to the number of specific DNA binding sites available.^17^ Therefore, the signal strengths of DNA-bound dyes cannot significantly differentiate unspecific binding sites, from specific binding with bacterial DNA, and are often similar in shape and size to the bacteria-soil aggregates leading to false positive identification of microbes.^15^ Therefore, bacterial counts of environmental samples will not be accurate with non-specific staining of DNA.^15, 17, 18^ Thus, for an accurate identification and counting, bacteria are usually detached^15^ for molecular biological and physio-chemical procedures to identify different organisms in their abiotic environments. As soil environments are disrupted for the extraction of nucleic acids or proteins,^4, 19, 20^ this approach leads to loss of essential biological information, such as the morphology of the cells, their spatial organization in soil, association with other organisms e.g. aggregates, biofilms in the natural environment, and individual cellular activities and functions.^1^ Therefore, these techniques cannot be used in a correlative microscopic approach to explore the global relationships between roots, microorganisms and chemical elements in the rhizosphere. CARD-FISH method can overcome issues associated with non-specific binding and improved visualization of bacterial cells in embedded samples.^21, 22^ However, the widely used resins for soil embedding including Araldite, Epon, and Lowicryl are not compatible for hybridization steps in CARD-FISH as media is hydrophobic once hardened, and thereby limiting the probe penetration and therefore clear identification of bacteria.^23^ Therefore, simultaneous characterization of microbes and correlation with the physio-chemical data within the intact rhizosphere soil are limiting.^21^ Thereby an embedding method allowing successful microbial identification *in-situ* and simultaneous high-resolution physio-chemical characterization is urgently needed for comprehensive understanding of microbe-soil-root functions.

A combination of 2D chemical spectroscopy, and microscopy, give different, yet, validating new information on the chemistry and organization of rhizosphere at biologically relevant scales. Thereby, such a methodological combination may be used to uncover and quantify biogeochemical processes in an intact rhizosphere with spatial resolution at micro-to nanometer scale.^11^ For high-resolution chemical microscopy, the soil needs to be dehydrated and embedded in various matrixes as a preprocessing step to preserve and stabilize its structure for high vacuum analysis.^11, 14, 17, 24-26^ Depending on the characterization technique, the embedding matrix can be gelatin,^27^ sulfur,^28, 29^ water glass^19, 30^ or a polyester, epoxy, or acrylic resins.^26, 31^ Among these, Araldite 502 is widely used epoxy resin for soil embedding due to its vacuum compatibility and fast outgassing property.^25, 26^ Araldite being a hard resin once solidified, can be cut and polished to achieve a smoother surface suitable for chemical microscopy, however, not compatible with immunohistochemistry methods or CARD-FISH. HIM, SEM-EDX, ToF-SIMS, nanoSIMS, confocal Raman spectroscopy (µ-Raman), and fluorescence microscopy, are promising for comprehensive characterization of specific chemical and microbial interactions within the rhizosphere. Capabilities of these instruments related to soil research is briefed in the SI section.

The correlation of multiple imaging techniques has received significant attention in recent years.^4, 11, 23, 31-35^ Using various techniques to characterize a single region of interest (RoI) (e.g. hot spot of microbial activity) within larger spatial context position can overcome each technique’s limitations leading to the understanding of multiple processes within the rhizosphere. However, sample preparation for a comprehensive correlative microscopic approach to study root, soil and spatial distribution of bacteria is a challenging task and demands compatibility with molecular biological methods, stability under high vacuum (10^−8^ NM^-2^), superior resin infiltration, minimal artefacts to the sample, and minimal topography of the analysis surface are to name few.

The focus of this work was to develop an embedding method which facilitates identification of bacteria and at the same time analyze the chemical composition of an undisturbed rhizosphere by means of correlative microscopy. However, preparation of a soil sample for various 2D and 3D characterization techniques with minimal destruction to the rhizosphere is a challenging task due to the sample size and the heterogeneous composition of organic and inorganic components. High spatial resolution techniques require mechanically stable and absolutely dry samples that are compatible with the required (ultra) high vacuum conditions. In this study, we developed a sample preparation method and a correlative image analysis workflow for soil sub-samples to comprehensively characterize spatial organization of roots, minerals and bacteria within *Zea mays* rhizosphere. Our approach consists of sample fixation, dehydration followed by impregnation with a vacuum compatible resin and molecular biology methods, sectioning with a waterjet to avoid polishing of the surface, finally imaging with multiple microscopic and spectroscopic techniques and analysis of results correlatively. To the best of our knowledge, microbial identification at the rhizosphere by CARD-FISH in a correlative high-resolution chemical microscopy workflow is a unique feature of our approach.

## Materials and Methods

### 2.1 Sample preparation

#### Soil Column preparation and subsampling

The soil substrate used is loam textured Haplic Phaeozem soil collected at Schladebach, Germany. Final soil column substrate corresponds to a bulk density of 1.26 g cm^-3^ (pH 6.21, sand 33.2%, silt 47.7%, clay 19.1%). Concise methodology on preparation of soil columns,^4^ substrate properties, and detailed subsampling method is documented elsewhere.^35^ Briefly, from the soil columns, 1.6 cm thick layers are cut at three different depths (5, 10, 15 cm) using a sharp blade without disturbing the soil structure (Figure 1a). These layers are carved into small metal cylinders (16 mm diameter). To prevent falling off of the soil top and bottom of these sub-samples were covered with a mesh and tightened with cable ties. In doing so, five sub-samples per depth were extracted out of which one cylinder was selected for resin embedding to prepare for 2D chemical and microbial imaging techniques. One from these sub-samples were then fixed with either 2% paraformaldehyde or Karnovsky solution.^36, 37^ Samples were rinsed twice with 1% PBS solution and stored at 4 °C in a 1% PBS solution until resin embedding. Remaining (so far unused) sub-samples were assigned to gain information on the gene expression and microbiome composition data derived from the same depth allowing joint interpretation of data across different disciplines in the future.

**Figure 1:**
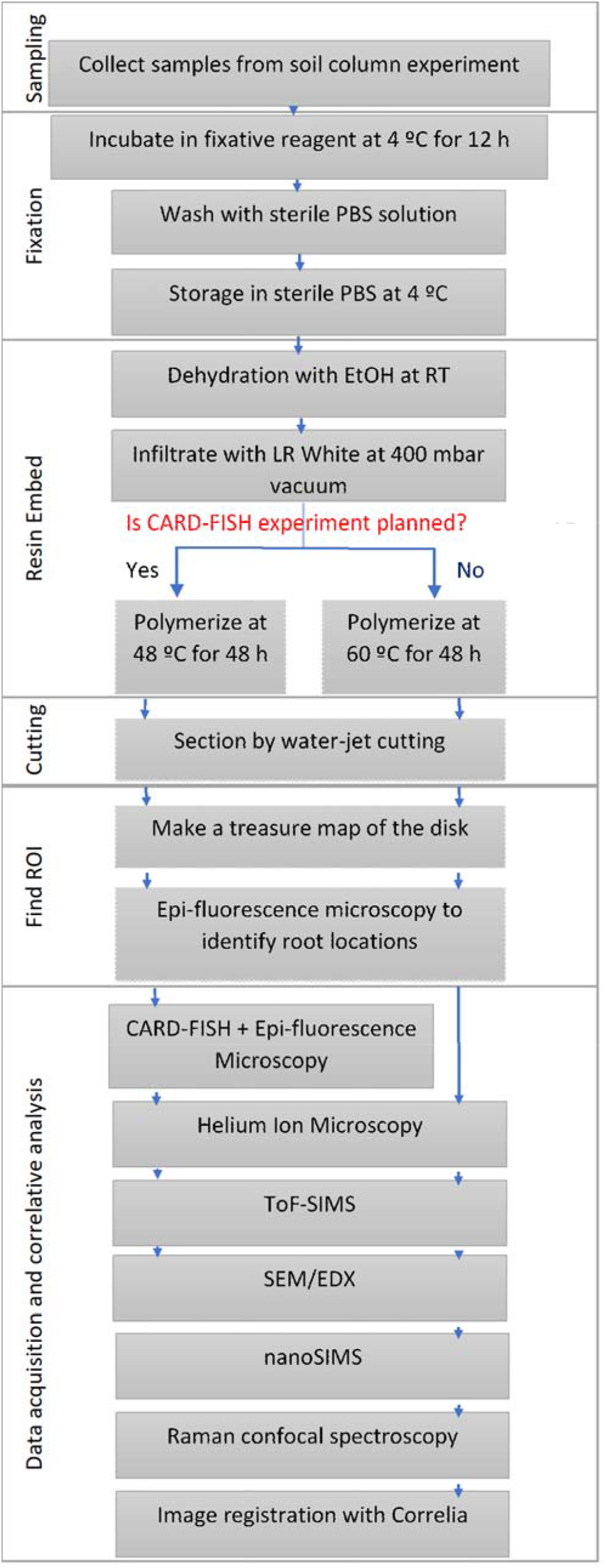
Schematic representation of the sample preparatio3n6, image registration and correlative analysis

#### Resin Embedding

Fixed samples were initially washed twice with 25 mL miliQ water by immersing in for 30 minutes each. The washed samples were placed in a 50 mL Eppendorf tube filled to 5 mL level with 2.85-3.45 mm size glass beads (Carl Roth GmbH, Germany) and dehydrated by immersing in increasing ethanol (Sigma-Aldrich) concentrations of 30%, 50%, 70%, 80%, 90%, 95%, and 99% for one hour each. Further on, samples were subjected to 100% ethanol (water-free) in three consecutive steps: 1) 1 h; 2) transferred to a new 50 mL Eppendorf tube (filled with fresh 2.85-3.45 mm size glass beads up to 10 mL level) and slowly pumped till 400 mbar vacuum and left for 14-16 h (overnight); 3) and immersed in a fresh 100% ethanol for 1 h before resin infiltration step.

London Resin (medium) (LR White) (Agar Scientific, UK) infiltration was carried out with consecutive steps of increasing EtOH:Resin mixture (vol/vol) 3:1, 3:2, 1:1, 2:3, 1:3, resin 1h each step. Next, the samples were treated with 100% resin three times for 1 h each. The sample was then transferred to a fresh 50 mL Eppendorf tube, filled with glass beads (to 5 mL level) and then filled up to 30 mL level with LR white resin, and kept for 14-16 h (overnight) under vacuum at 400 mbar, with continuous purging with Ar gas. Care was taken to slowly reduce the vacuum to 400 mbar. Finally, tubes were exposed to Ar gas for 2 minutes, tightly sealed and samples cured in a water bath for 48 h at either 48 °C or 60 °C.

The sample was dehydrated with ethanol and then substituted gradually with resin by increasing resin concentrations. Gradual dehydration and infiltration reduce the potential of causing any distortion to ultrafine cellular structures and the microbial colonies and less lipid component are removed.^1^ We have used ethanol instead of acetone as this is weaker solvent compared to acetone and therefore less prone to extract cell components such as lipids. Glass beads were used during dehydration and embedding steps to lift the sample from the surface and keep it straight during the process. This stops air trapping at bottom and helps exchanges from the top and bottom surfaces to increase the embedding efficiency. This also helps positioning sample during the water-jet cutting. Care was taken at every step to make sure the transfer of solvents take place slowly and to prevent air trapping in between these glass beads, which can result in poor infiltration.

#### Water-Jet cutting

The cured resin embedded samples were securely mounted in a horizontal plane on to the waterjet cutting system (Premium Cut 3D; StM Waterjet GmbH, Salzburg, Austria) using an ad-hock designed clamp. A combination of pressurized water and abrasive garnet particles cut the sample at predefined positions. Initially, abrasive particles of two mesh sizes (Mesh-50 of 300 µm in diameter and Mesh-200 of 75 µm diameter) and different settings are used to optimize the final surface roughness using pure resin. These settings and resulted roughness are listed in the SI section. The embedded soil sample was cut with the settings that resulted in minimum surface roughness. Care was taken not to cut the soil sub-sample completely to prevent it falling into water bath during the cutting procedure. Once multiple cuttings were performed and dismounted from the instrument, samples were washed three times by immersing in MiliQ water for 10 minutes each. Individual disks (1 mm thick) were cracked to separate and air-dried before stored in a gel –box (Gel-Pak^®^) until further analysis.

#### CARD-FISH

The LR White embedded soil disks were divided into multiple pieces for hybridization using a mixture of the following probes: EUB 338I, II, and III,^38, 39^ to target a wide range of soil bacteria in the sample. This protocol follows Pernthaler *et al*.^*40*^ with following modifications. Embedded soil disks were permeabilized in a Lysozyme solution (10 mg/mL in 0.05 M EDTA, pH 8.0; 0.1 M TrisHCl, pH 7.5) for 1h at 37 °C, followed by washing with ultrapure (miliQ) water and further treatment with Achromopeptidase for 30 min at 37 °C. The enzymatic permeabilization was followed by 3x washing with miliQ water and treated for inactivation of endogenous peroxidases in 0.15% H_2_O_2_ in methanol (vol/vol)) for 30 min., and further on followed by washing 3x with ultrapure water at room temperature (RT) and dehydrated by increasing EtOH series 50%, 70% and 96%, 30s each and air dried. The hybridization was done overnight (app. 20h) at 35 °C in a hybridization buffer containing (0.9 M NaCl, 40 mM TrisHCl, 10% (w/v) dextran sulfate, 0.01% (w/v) sodium dodecyl sulfate (SDS), 10% blocking reagent, 1x Denhard’s reagent, 0.26 mg mL^-1^ sheared salmon sperm DNA, 0.2 mg mL^-1^ yeast RNA, 5M NaCl 3.6 mL, and 50% Formamide. The HRP-labelled probes (Biomers, Ulm, Germany) applied are specific for bacteria were used at a concentration of 0.166 ng mL^-1^ (HRP-probe stock solution of 50 ng mL^-1^ diluted 1:300 v/v in hybridization buffer). Following hybridization, resin embedded soil pieces were incubated in 50 mL of pre-warmed washing buffer containing 19mM NaCl, 5mM EDTA (pH=8.0), 20mM Tris–HCl (pH=7.5), and 0.01% SDS for 15 min at 37°C. After washing, samples were incubated for 15 min at RT in 1× PBS (pH=7.6) to equilibrate the HRP-labeled probe. Subsequently, tyramide deposition was performed by incubation for 30 min at 46°C in the dark in amplification buffer containing 1× PBS, 2M NaCl, 0.1% (w/v) blocking reagent, 10% (w/v) dextran sulfate, 0.0015% (v/v) H_2_O_2_, and 1 µg mL^-1^ Alexa 594-labeled tyramides (ThermoFisherScientific, Waltham, Massachusetts,USA). Afterwards, samples were rinsed in 1× PBS for 10 min at RT followed by counterstaining with 4’,6-diamidino-2-phenylindole (DAPI) 1µg mL^-1^ for 10 min at RT, washing in ultrapure water and air dried. Samples stored at -20 °C until imaging.

### 2.2 Correlative Imaging

#### Light Microscopy

The embedded and cross-sectioned soil disk was mapped entirely with reflected light microscopy, using a WITec alpha 300 confocal Raman microscope (WITec GmbH, Ulm, Germany) equipped with the true surface® option to make an overview montage. The acquired 2704 individual mosaic images with 136 stack layers were stitched in auto mode with continuous movement algorithm by WITec project five (5.2) software to make a single figure of 15.857 mm x 15.857 mm. This overview is the treasure map to locate region of interests (RoI) during various microscopy analysis. As roots could not be efficiently located by this method, epifluorescence imaging was adapted to find areas with roots (RoI) for further analysis for other imaging techniques.

#### Epifluorescence Microscopy

To find the roots embedded in the soil, resulted disks were scanned with epifluorescence microscopy (Zeiss AxioImager.Z2) equipped with an HXP R 120W/45C UV Hg-vapor lamp, Colibri.2 LED illuminations (590 nm, 470 nm and 365 nm) and imaged with objective lens 20x (EC Plan-Neofluar/0.50 M27). Fluorescence DAPI filter set was used to visualize autofluorescence of the root. Images were acquired with a color CCD camera connected to an imaging software (Zeiss AxioVision).

CARD-FISH hybridized samples were imaged with fluorescent filter sets (DAPI (365 nm), DsRed (590 nm)), using an 20X (NA=0.50) and 50X air objectives (NA=0.55) and a black and white CCD camera (AC MR R3) connected to Zeiss AxioVision. Acquired images were false colored to respective channel, DAPI in blue and DsRed in orange.

#### Helium Ion Microscopy (HIM)

After epifluorescence microscopy mapping regions-of-interest were imaged with a scanning helium ion microscope (Zeiss Orion NanoFab, USA) The reason for using HIM instead of SEM is that this surface-sensitive method can be considered almost non-destructive: Neither a significant damage by the ion beam has to be expected (only about 30 He^+^-ions are implanted per pixel) nor does it require a metal- or carbon-coating of the surface, as is needed for SEM to avoid charging. HIM imaging was carried out with He^+^ at 25 kV acceleration voltage and a beam current of 0.1 to 0.3 pA. Secondary electron images were collected in a quadratic field-of-view of 300 µm. The dwell time amounted to 0.5 µs and 32 times line averaging was done to reduce noise. For charge compensation an electron flood gun was used during imaging. In total nine fields-of-view were acquired with 25% overlap and stitched together using either the pairwise stitching^41^ or mosaicJ^42^ plugins in Fiji to create a larger map of the RoI. The obtained HIM microscopies of the region-of-interests are rich in features and thus serve as an ideal base for the registration of other modalities.

#### ToF-SIMS experiment

The resin-embedded rhizosphere sample was analyzed with a TOF-SIMS 5 instrument (IONTOF GmbH, Münster) employing 30 keV Bi_3_^+^ cluster ions (LMIG NanoProbe source) as primary projectiles. Secondary ions were analyzed in negative and positive extraction modes. With a 100 µs period of primary pulse repetition, the sample surface was exposed to 0.05 pA of Bi_3_ ^+^ probe and the secondary ion species were detected within 12-1200 Da range in Delayed-Extraction mode allowing for 120-250 nm lateral resolution and about 5000 MRP.^43^ Analysis of 250×250 µm^2^ size RoI’s were performed by rastering the Bi_3_ ^+^ probe randomly in 2048×2048 px raster-pattern reaching 2 shots per pixel in 28 minutes of single analysis frame. After acquisition of each analysis frame, an area of 500 µm x500 µm or 800 µm x800 µm around the analysis crater was sputtered with either i) 75 nA of 1 keV Cs^+^ or ii) 250 nA of 1 keV O_2_ ^+^ beam in 200-400 frames for 3-7 minutes to analyze the complex heterogeneous sample in depth. Sputtering with Cs^+^ ion beam was examined in negative as well as positive extraction mode, whereas sputtering with O_2_ ^+^ was examined only in the positive extraction mode. To compensate for surface charging of insulating uncoated samples, flooding with Ar gas (partial pressure 2×10^−6^ mbar in analysis chamber) was employed in combination with 21 eV electrons from pulsed neutralizing electron gun (NEG) during the analysis. Between 80 and 200 planes were acquired per analysis. The lateral drift, mass-shift corrections and further data evaluation steps were performed with the Surface Lab software (Version 7.1, ION-TOF GmbH, Germany) to generate peak lists and corresponding ion distribution maps. Negative mass spectra were calibrated using the mass peaks assigned to H^-^, C^-^, CH^-^ O^-^ and OH^-^. The peak lists were generated by the automatic peak search program between 0-1000 m/z mass range. This step has resulted 687 mass peaks for multivariate analysis. Initially, PCA was done to determine the number of principal components for the analysis. Though 155 PCs were suggested for this sample, we have only selected 10 PCs for further analysis in MCA to determine their corresponding spectra.

#### NanoSIMS experiment

After ToF-SIMS experiments, the resin-embedded rhizosphere samples were mounted on sample holder, 5 connecting bridges were made for better conductivity (see SI) and coated with 20 nm Au/Pd (80/20%) layer to provide a conductive surface and analyzed with a nanoSIMS 50L instrument (CAMECA, AMETEK) in negative and positive extraction modes.

In the negative extraction mode, 16 keV Cs+ ions with a beam current of 3 pA beam were employed to analyze 95×95 µm^2^ areas in sawtooth raster of 512×512 pixels. The dwell time was 2 ms/pixel and a field aperture of 300 µm diameter was used. Prior to the analysis, the targeted field of view (FoV) of 150 µm x150 µm was pre-implanted with 200 pA Cs+ beam for 20 minutes. Mass Resolving Power (MRP=M/dM) above 7000 was achieved with 20 µm ×140 µm (width × height) entrance slit, 200 µm ×200 µm aperture, 40×1800 µm exit slits and the energy slit cutting-off 20% of ions at their high-energy distribution site, for the collected secondary ion species (^12^C_2_^-, 13^C^12^C^-, 12^C^14^N^-, 13^C^14^N^-, 31^P^-, 32^S^-^, and ^31^P^16^O_2_-).

In the positive extraction mode, 5 nA O^-^ beam was pre-implanted in an area of 150 µm x150 µm for 30 minutes. The analysis has been implemented with 20 pA of 16 keV O^-^ ion beam in 100×100 µm^2^ sample areas (within those preimplanted) in 512×512 pixels with dwelling time of 2 ms/pixel and the field aperture of 300 µm diameter. Mass Resolving Power (MRP=M/dM) above 6000 was achieved with a 20 µm ×140 µm (width × height) entrance slit, 350×250 µm aperture, 40×1800 µm exit slits and the energy slit cutting-off 10% of ions at their high-energy distribution site, for the collected secondary ion species (^11^B^+, 24^Mg^+, 27^Al^+, 31^P^+, 40^Ca^+, 55^Mn^+^, and ^64^Zn^+^).

The acquired 30-100 plains per analysis were corrected for lateral drift and accumulated using the Look@NanoSIMS software (LANS).^44^ Individual data files were further processed by the OPENMIMS ImageJ plugin to extract only 5 image planes and to create a combined field of view showing larger area of rhizosphere using mosaic J plugin ^42^ in Fiji.

#### SEM-BSE and SEM-EDX

In order to obtain mineralogical information about the RoI selected based on the fluorescence micrograph, the surface was analyzed with a field-emission SEM (Zeiss Merlin VP Compact) in BSE-mode and coupled to an EDX spectrometer (Bruker Quantax XFlash 5060F), respectively, at an electron acceleration voltage of 11.8 kV and a beam current of about 250 pA. Firstly, the surface was scanned using the in-lens electron detector negatively biased at 958V to completely suppress secondary electrons but allow high-energy back-scattered electrons to be detected. This resulted in high-resolution images of the rhizosphere providing a material contrast sufficient to differentiate minerals, root cells and resin infiltrations. To obtain more specific information about the elemental composition SEM-EDX maps were collected using a Bruker Quantax XFlash 5060F energy-dispersive X-ray spectrometer (Bruker Nano GmbH Berlin, Germany). The resulting element-distribution maps were background-corrected by removing the bremsstrahlung using a physical model implemented in the software Bruker Esprit 1.9. The SEM-BSE as well as the SEM-EDX maps are conveniently registered onto the HIM micrographs (for SEM-EDX using the Si channel for registration of the entire set) employing automatic rigid mutual-information method implemented in the ImageJ plug-in Correlia.^45^

#### Confocal Raman Spectroscopy

Selected root-soil interfaces were analyzed with the same WITec alpha 300 confocal Raman microscope (WITec GmbH, Ulm, Germany) used to acquire the overview light micrograph serving as treasure map. For excitation a solid-state laser emitting in the infrared (785 nm) was used. The analysis was carried out at a laser power of 0.5 mW and spectra were acquired with a grating monochromator (600 g/mm) and CCD camera detection. Mapping was done in an area of 160 µm x 160 µm, with 0.25µm pixel-size. The objective lens was a Zeiss LD plan-NEOFLUAR 20× air objective lens with 0.4 numerical aperture, allowing resolution of less than 1µm. The WITec project five (5.2) software was utilized for the data processing. After cosmic ray removal and background subtraction, the automated statistical evaluation of all spectra by *k*-means cluster analysis resulted in spectral unmixed images which provided an insight into the chemical composition of the sample.

Soil minerals were analyzed with the same system but using 532 nm wavelength and 33 mW laser power instead. An area of 170 µm x 170 µm was scanned with an integration time of 10 s and 85 point per x,y direction of scan resulting 861 s per line scan using the 20× air objective lens.

## Results and Discussion

Soil samples for high-resolution microscopy must be dry, stable, tolerant of ultra-high vacuum (10^−8^-10^−10^Torr) and conductive unless charge compensation is not available. As such water containing samples like soil needs to be pre-processed before characterization. In general, it is also expected to be flat and highly polished to acquire the best characterization possible. Therefore, soil sample preparation for high-resolution microscopy involves stabilizing biological components (fixation), removing water (dehydration), resin-embedding, and polishing.^14^

Fixation, dehydration and resin embedding are important initial steps in preserving the original morphology of living microbial cells and plant roots in their original condition.^1, 46^ Therefore, we have performed fixation immediately after collecting the samples from the columns to limit structural alterations that may occur to roots and microbes during sample preparation and storage. Considering the complexity and size of the rhizosphere samples, the high pressure freezing and freeze substitution methods are not compatible.^14^ Therefore, the Karnovsky fixative^36, 37^ was adapted in our sample preparation. Aldehyde-based fixatives preserves cells by crosslinking proteins, which results in the stabilization of the cellular fine structure. It is crucial to select the right fixation method when chemical characterization of the sample is planned. Several studies show that the choice of fixative and their time of action can affect the overall chemistry of the sample.^46-49^ Widely used fixatives such as glutaraldehyde and ethanol can cause significant changes to the Raman spectra of bacteria indicating chemical alterations, whereas formaldehyde and sodiumazide were better at preserving spectral features.^48^ As rhizosphere consists of soil, root, bacteria and other organic and inorganic material, various components can be studied, and it is not trivial to adapt a single fixative method. We have used ethanol instead of acetone, to minimize the membrane damages by acetone. Another critical factor to be considered is the choice of resin as this could change the chemistry of the sample.^50^ The ‘*ideal*’ resin may only introduce minimal physiological and chemical artifacts to the test sample, and it has to be compatible for multiple imaging requirements of various analysis methods. Overall, it is important to note that the fixation and resin embedding steps have to be carefully selected depending on the specific analysis planned. The soft LR white resin we choose is used to embed and preserve microbial cells in various tissues, the environmental samples in correlation with FISH and TEM to study functions of microbes in natural systems^22, 23, 51^. However, for the best of our knowledge, LR white has not yet been used for soils or rhizosphere samples in high resolution workflows where bacteria-plant-organic-inorganic interphases are correlatively characterized.

The Helium ion micrographs in Figure 2c demonstrate the quality of embedding with LR white using this procedure. Overall, the quality of infiltration is satisfactory as: neither holes representing poorly infiltrated areas nor cracks representing changes in the soil structure are noticed in the resulted samples after water-jet cutting. There were no gaps found at the root-soil interface, and minerals are in contact with the outer root wall and minerals of different sizes are held by the cured resin. The root cells in the center of the image are well infiltrated by resin without vicinity of pores, holes or non-cured areas. HIM images mainly show a topography contrast with pronounced edges. The weak contrast in Figure 2c is an indication for the low number of edges noticeable, indicating consistent infiltration of rhizosphere. Consistency in resin infiltration and curring are mainly due to the low viscosity of LR white compared to common polyester (380-800 mPas) and epoxy (200 mPas) resins. This allows LR white resin to effectively penetrate through the soil pore network and into root cells to preserve rhizosphere for microscopic analysis in (ultra-) high vacuum conditions.^52^ However, within the root, sub-micron scale space between cell wall and cell cytoplasm is evident. This is due to the shrinkage of resin during the curing process. This separated space is also apparent between the soil and the Al cylinder of the soil-sample (data not shown). However, LR white shows the minimum shrinkage of 2% during impregnation and curing compared to most commonly used epoxy and polyester resins, which showed 6.4% and 7.5% shrinkage, respectively.^52^ LR white is sensitive to oxygen, therefore, samples were processed in a desiccator purged with Ar during infiltration and sealed when curing. More often, traditional soil-related embedding demands hard-plastic resin to polish the surface, where soft LR white resin does not fit to prevent the smearing during cutting and polishing steps.^52^ Softness can lead to excessive smearing and delocalization compared to hard resins during cutting and polishing. Consequently, we have adapted water-jet cutting procedure to slice the embedded soil and avoided polishing.

**Figure 2:**
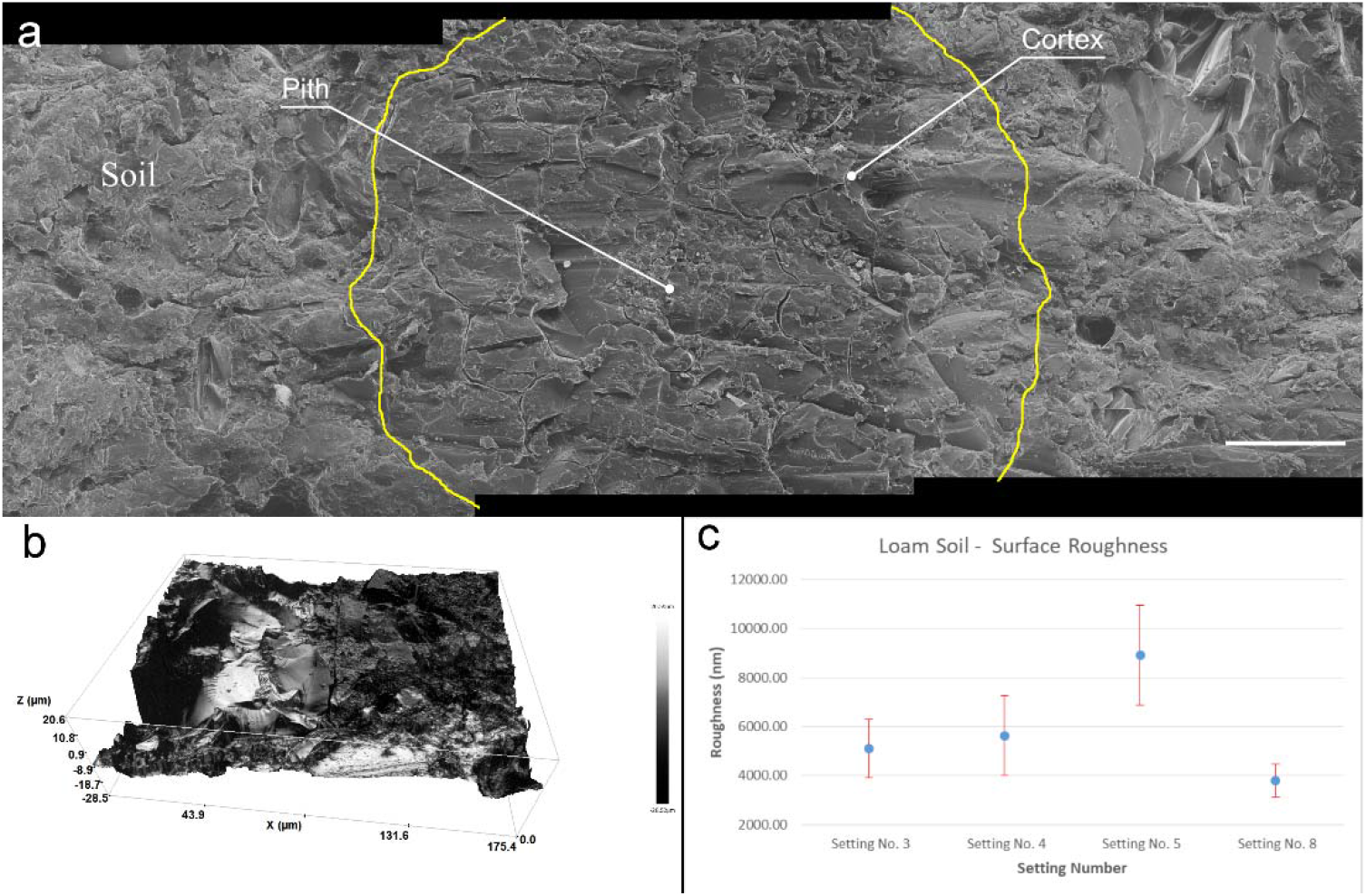
Surface roughness of embedded soil sample. a) Three Helium ion micrographs are stitched showing the quality of embedding and surface roughness of a root after water jet cutting. Root cell is located in the middle image and outlined in yellow for easy recognition. Root cells and soil-mineral interface are well infiltrated by the embedding procedure. Scale bar 50 µm b) Surface profile of a root area measured by profilometer before microscopy. Rz= 49.8 µm c) Resulted surface roughness of embedded soil once cut with different parameters by water-jet.

### 3.1 Water-Jet cut and Surface Roughness

As the RoI of the rhizosphere is at the microscale, variations in the surface roughness might affect the nanoSIMS and ToF-SIMS measurements. To achieve a minimum surface roughness, we have optimized parameters for cutting the resulting surface; roughness was measured (See SI) to achieve the minimum roughness of 40.0±5.00 nm (Figure 2b). Figure 2b shows the resulted 3D roughness profile of the area where we have carried out correlative microscopic analysis. However, we have noticed roughness of embedded soil is random and highly changes locally compared to pure resin. Our results demonstrates that an Rz value of approximately 50 nm (Figure 2a) can provide comprehensive chemical measurements of the rhizosphere. Moreover, water-jet approach can be used to gain multiple disks with 1 mm thickness (See SI) that allow to gain multiple cross-sections from single sub-soil sample. Thereby if need arise, can characterize multiple planes, which is an advantage for comprehensive understanding of the chemical gradients within the rhizosphere.

Water-jet is a versatile cutting technique adopted in various industries to cut heat-sensitive samples without resulting rough, barring edges. As an additional advantage, water-jet cut avoids heat generation during the cutting. Thereby, avoid possible heat damage for bacterial ribosomes, which might affect the CARD-FISH experiment. Further, this process only uses water and sand and does not produce any toxic gases during processing. Thereby environmentally friendly and can be directly discharged.

Surface imperfections on the resulted surface scatters light and decrease the quality of optical images of large fields of view. Therefore, polishing is a regular step adapted to prepare soil samples to achieve quality of visual images with fields of views of mm to cm scales. Yet to achieve flat surfaces without scratches or furrows are difficult with samples like soil with a mixture of components of varying hardness. Polishing can also increase surface furrows and deterioration of soft crystals, remove attached bacterial cells on the minerals, smearing, and unknown manipulation of the sample at the micro to nano scale. Therefore, we avoided the polishing of the surface, thereby, surface smearing and dislocation of materials at microscales by adapting water-jet cutting during soil sample preparation.

### 3.2 Selecting Region of interest (Root/ Bacteria) by Light microscopy

Bright- and darkfield reflective images resolved resin from mineral particles but cannot be used to identify roots or bacterial colonies in the rhizosphere (Figure 3). Homogeneous resin infiltration makes it challenging to select the region of interest, which is a root in this study. Therefore, fluorescence microscopy is used to locate the region of interest (RoI) for further high-resolution microscopy. Epifluorescence microscopy under UV captures auto fluorescence of embedded roots (cyan), and resin (dark blue), and minerals in dark color (Figure 3, inset), differentiating them from each other. The dark blue colored resin is the soil matrix’s air pores, now infiltrated by resin in this procedure. The selected area for further analysis is well focused, while a dominant scratch and bottom section of brightfield and fluorescence images are blurred. As the sample is not polished, and the depth of focus is out of the objective lens focus range, blurred sections appear. However, the root and the surrounding microscale area is in focus and is sufficient to obtain a clear image. Therefore, avoiding of polishing step has not influenced the image quality within RoI. There is a prominent tan colored border that is visible in some parts around the root and minerals. These correspond to Ca and P in EDX and nanoSIMS micrographs (Figure 3).

**Figure 3:**
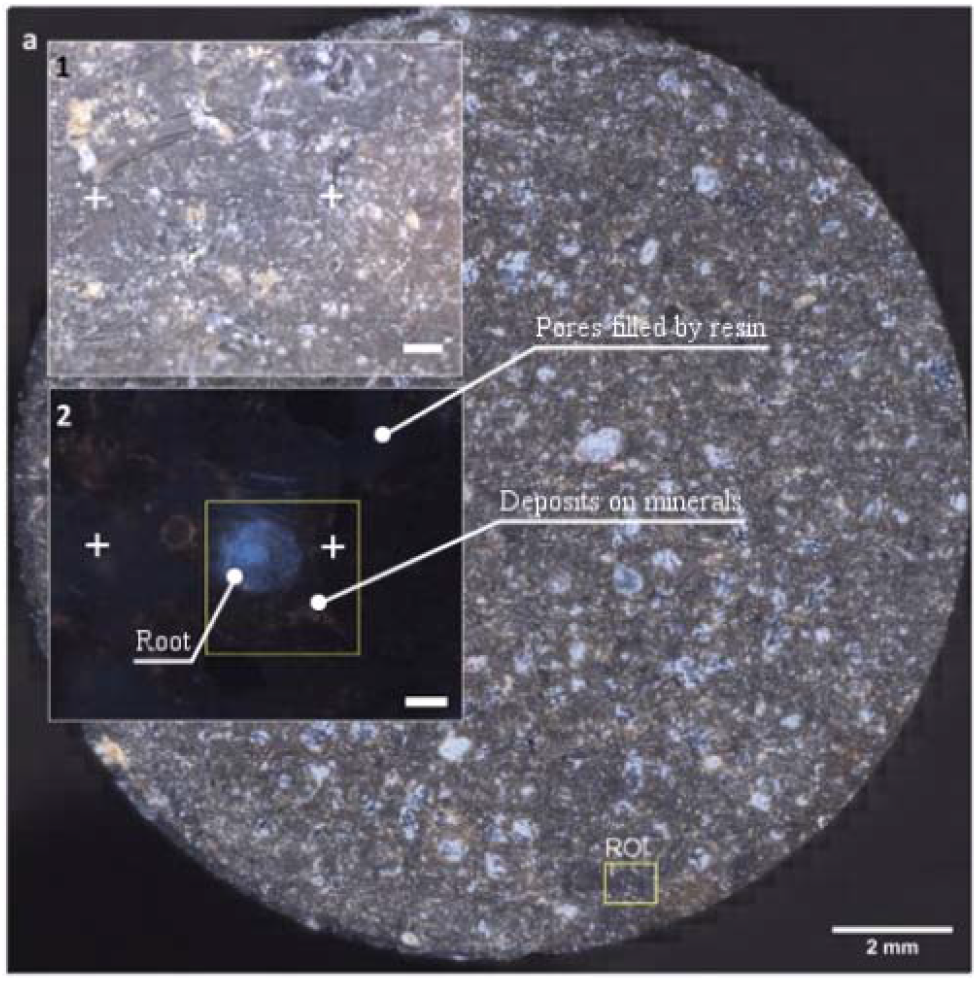
Resulted disc of soil column after water-jet cutting. Disk is a stitched image from 2704 individual images at darkfield microscopy mode. Insets 1-2 shows the magnified RoI marked in the soil disk. Crosses are surface features used to identify the RoI under different microscopes. 2) Epi-fluorescence micrograph of RoI showing the root, soil and minerals. Square marked in yellow is the RoI for high-resolution microscopy under HIM, ToF-SIMS, SEM-EDX, nanoSIMS and µ-Raman.

### 3.2 Chemical Microscopy

#### ToF-SIMS

We could identify the resin, minerals and the root clearly using the negative ion extraction mode analyzed by PCA-MLS (Figure 4). Soil matrix was identified by the C_x_H_y_O_z_^-^ and SiCH_4_O_2_^-^ fragments and the cell walls of the root are identified by CN^-^ and CNO^-^fragments. There are CNO^-^ fragments scattered randomly within soil matrix, which are indicated as green color tiny spots in the Figure 4. These may correspond to plant exudates or organic matter within the soil matrix. A pore area immediately above the root is filled with resin and appear separate to the root and soil matrix. LR white resin is composed of C, H, and O (See SI for EDX spectra) and therefore, resin related fragments can be distinguished by the C^-^(m/z=12), CH^-^(m/z=13), C_2_^-^(m/z=24), C_2_H^-^(m/z=25), CH_2_^-^ and O^-^ fragments (See SI). Given the availability of other unknown hydrocarbon molecules within the soil matrix, it is difficult to distinguish resin C_x_H_y_O_z_^-^ fragments from the organic molecules which may be already present within soil matrix using currently available PCA methods. Nevertheless, with PCA we could easily distinguish root, PO_x_^-^ and minerals. However, with future improvements in the complex analysis methods, and with an availability of a correlative sample preparation method, distinguishing biomolecules from LR white resin may be possible.^9^

**Figure 4:**
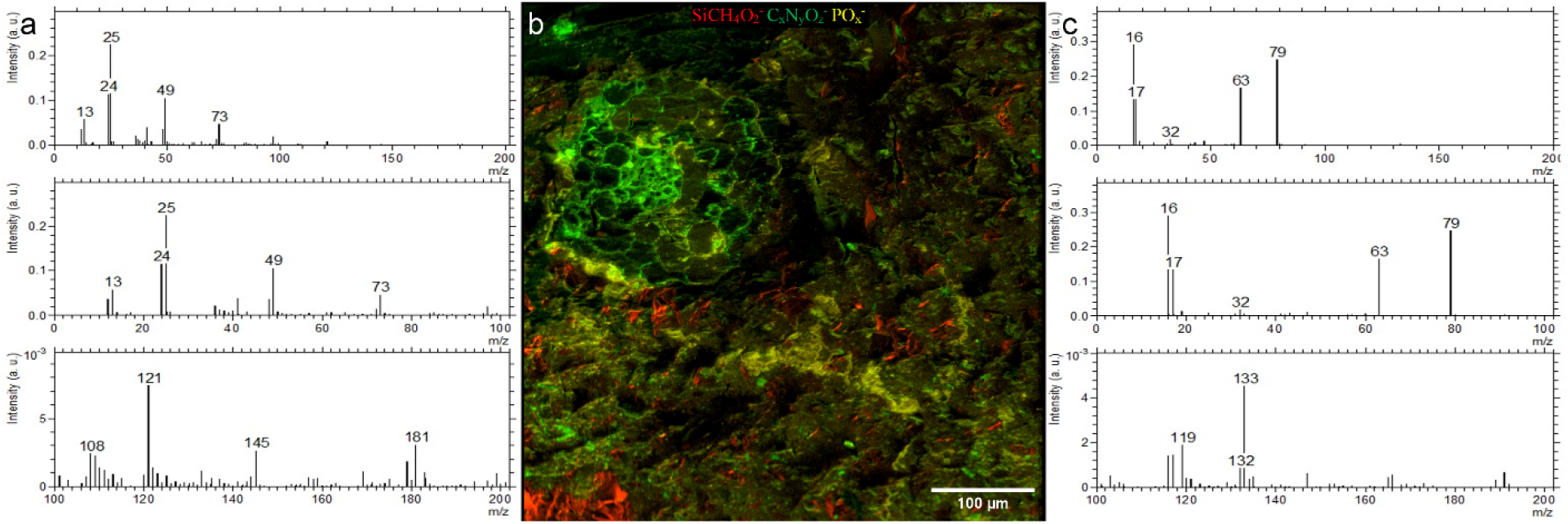
ToF-SIMS analysis of a Rhizosphere embedded sample. a) Spectra of LR white b) Molecular fragment distribution related to SiCH_4_O_2_^-^, CNO^-^ and PO_x_^-^ c) Spectra of PO_x_^-^ rich areas.

On the root surface PO_x_ ^-^ deposition was recognized, on one side of the outer root wall, which spreads into the soil matrix in the analyzed plane, and correlates with nanoSIMS, EDX, and fluorescent image (UV) data. These precipitations may be related to the physio-chemical interactions between root’s nutrient uptake mechanisms.^53^ PO_x_ ^-^ precipitations were also noticed in soil matrix and are likely to occur as P anions highly associated with cations like Al^3+^, Fe^3+^ and Ca^2+^ and tends to firmly absorbs onto clay minerals and metallic oxides.^53-55 53-55^ Moreover, we have used the positive extraction mode to identify the different minerals and elements. (See SI). K-related, Si-related and Na-related minerals were easily distinguished by PCA. We show that, the positive extraction mode is complementary to EDX analysis and the topography resulted by water-jet cutting has minor influence on data acquisition. Compared to EDX, ToF-SIMS has a better sensitivity and spatial resolution, thereby combination of both positive and negative extraction modes in ToF-SIMS analysis can be used to study rhizosphere PO_x_ ^-^ distribution and and bioavailability of P at various stress conditions.

#### SEM-BSE and SEM-EDX

Both, SEM-BSE and SEM–EDX maps provide chemical information about the surface of soil-disk. Electrons back-scatter more efficiently on heavy than on light elements, such that SEM-BSE contrast is used to distinguish regions of low electron density, resin and soil-organic matter, from electron-dense ones, namely mineral particles as shown in figure 5. In this way mineral classes of different electron density are distinguishable, for instance sodium-feldspar and quartz from iron-minerals. In addition, element-distribution maps were measured by SEM-EDX. In principle any element heavier than Be can be measured by SEM-EDX with increasing sensitivity for heavier elements. Major element mapping was feasible starting with C such that, the concentration of B was too low to be detected with SEM-EDX. Both, SEM-BSE and SEM-EDX micrographs proved a good overview on the soil composition (Figure 5). Mainly Si, Na, and Al containing minerals, e.g. quartz and Na-feldspar, were identified but also Ca, P and Fe are detected. At the root wall Ca precipitation is noticed. This may be due to the nutrient uptake regulation mechanisms by the plant root or the changes in pH.^53-58^As LR white is composed of C, H, and O, the infiltrated resin is clearly identified as C in SEM-EDX maps. Using SEM-BSE, resin can be separated from the soil matrix as shown in Figure 5. It should be noted that LR-white contains traces of Na, Al, Si, S and Cl (See SI) which must not falsely be interpreted as minerals. Also, the separation and identification of soil organic matter by SEM-EDX is challenging once embedded in LR-white. Nevertheless, the correlation of HIM, SEM-BSE and SEM-EDX is an ideal tool to characterize morphology and elemental composition of minerals in soil.

**Figure 5:**
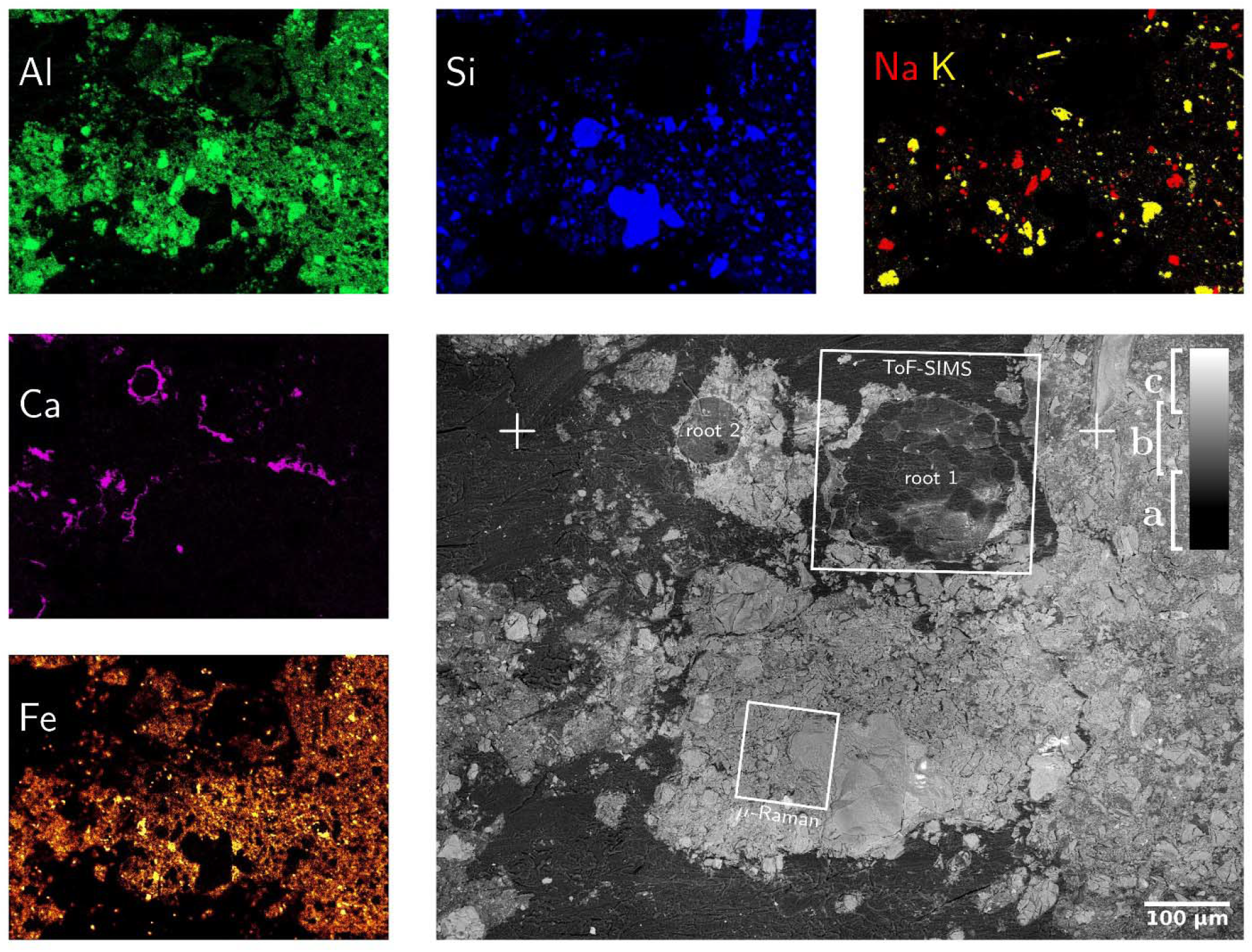
SEM-BSE (gray figure) and SEM-EDX maps of mineral-forming elements. SEM-BSE contrast allows for separation of resin (a), light (b) and heavy (c) minerals as well as the identification of root-cells. The positions marked as “+” are identical with those in figure 3. The SEM-EDX false-color maps show exactly the same field-of-view such that SEM-BSE contrast can be linked to elemental composition. Please note that the Calcium distribution is almost identical to that of phosphorous which hints to calcium phosphates precipitated around the root surface.

#### nanoSIMS

NanoSIMS negative mode imaging shows the distribution of ^12^C^14^N^-, 31^PO_x_^-^ and ^32^S^-^ secondary ions in the embedded rhizosphere sample. The ^12^C^14^N^-^ helped to differentiate the organic matter from the resin. (Figure 6 a-c). Therefore, it shows the cell arrangement of the root. ^31^PO_x_ ^-^ was mainly found at the outer root wall and extend to the soil matrix. At the side where ^31^P is precipitated on the root epidermis, ^31^P was not abundant in the cortex cell walls. On the side where ^31^P is not precipitated, ^31^P is distributed within cortex cell walls of the root (See SI). ^32^S^-^ was detected on cortex cell walls.

**Figure 6:**
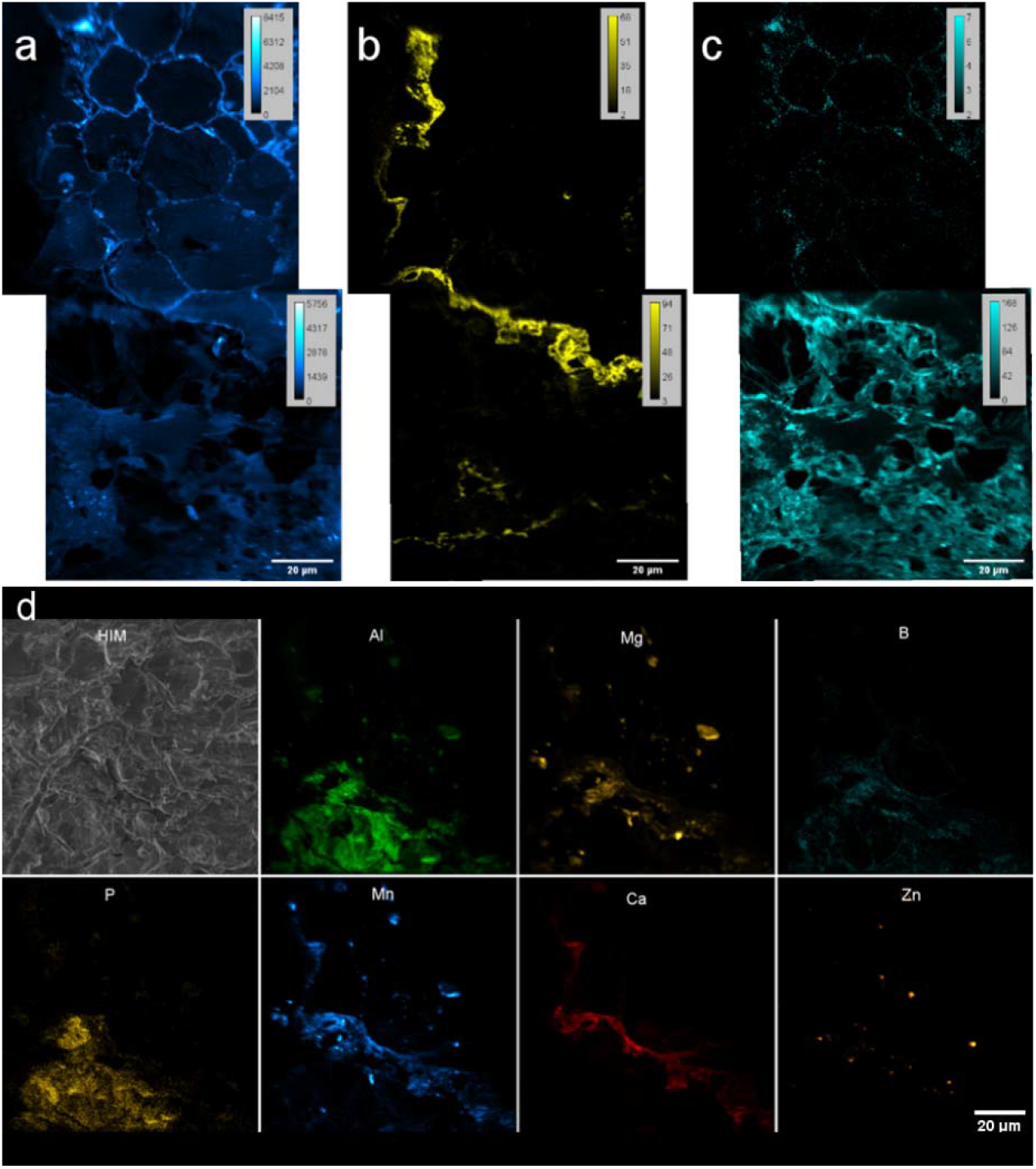
nanoSIMS images of resin embedded Rhizosphere. a-c) Spectra from negative extraction mode showing elemental distribution a)^12^C^14^N^-^ b) PO_x_^-^ and c) S^-^ at root-soil interface of rhizosphere. Two FoV’s stitched by MosaicJ in Fiji. d) Maps from positive extraction mode showing elemental distributions of ^27^Al^+, 24^Mg^+, 11^B^+, 31^P^+, 55^Mn^+, 40^ Ca^+^ and ^64^Zn^+^. d-HIM shows the HIM micrograph of analyzed area.

With the positive extraction mode, we have identified the distribution of ^27^Al^+, 24^Mg^+, 11^B^+, 31^P+, ^55^Mn^+, 40^Ca^+^ and ^64^Zn^+^ (Figure 6 d). Though Boron could not be identified by EDX, capability of detection with nanoSIMS allowed to study B in rhizosphere as an important micronutrient for the quality and yields of crops, and a cell wall component.

Combination of positive and negative extraction modes showed distribution patterns of PO_x_^-^, highly correlates with cations Ca^2+^ and Mn^2+^ and some to extent of Mg^2+^ and Al^3+^ at the root’s outer surface (Figure 6b, d-Ca). This can be related to the co-precipitation of P as highly insoluble Ca, and Al salts reducing the bio-availability of P for plants.^54-56, 58^ Possibility of detecting ^12^C^14^N^-^,^31^PO_x_^-, 32^S^-, 27^Al^+, 24^Mg^+, 11^B^+, 31^P+, ^55^Mn^+, 40^Ca^2+^ and ^64^Zn^2+^ allow to study the micro nutrient related research and their pathways within rhizosphere coupled to various correlative techniques.

#### µ-Raman

Employing WiTEC’s True Component analysis^®^ after µ-Raman mapping we could distinguish quartz minerals from Al-rich feldspar minerals similar to SEM-EDX. Figure 7a shows the map of quartz and its spectrum (Figure 7b), with the prominent peak at 468 cm^-1^ corresponding to Si-O-Si stretching. Figure 7c and Figure 7d shows an Al-containing mineral and its spectrum, respectively, identified in the same field-of-view. Here µ-Raman adds additional knowledge to SEM-EDX by revealing that the spectrum indicating measured minerals are a mixture of two minerals, which consists of 59.52% Zircon and 40.48% of ammonium and sulfate containing minerals rather than a single pure component (Figure 7d), whereas SEM only provided elemental information (See SI). The dual peaks at 350 cm^-1^ and 436 cm^-1^ and peak with a shoulder at 1000 cm^-1^ may corresponds to Zircon while the sharp peak at 995 cm^-1^ as well as two smaller yet broader peaks at 450 cm^-1^ and 600 cm^-1^ correspond to ammonium sulfate (see SI). Possible source would be aluminium sulfate and ammonium nitrate which were supplied as nutrients during the soil column preparation.^35^ The Raman spectra of LR white resin has a sharp peak with two shoulders at 2950 cm^- 1^ corresponding to C-H stretching and a small sharp peak at 1450 cm^-1^ corresponding to C=O (See SI). Therefore, using Raman spectroscopy we could accurately identify interested minerals in the resin imbedded sample, but not the biomolecules.

**Figure 7:**
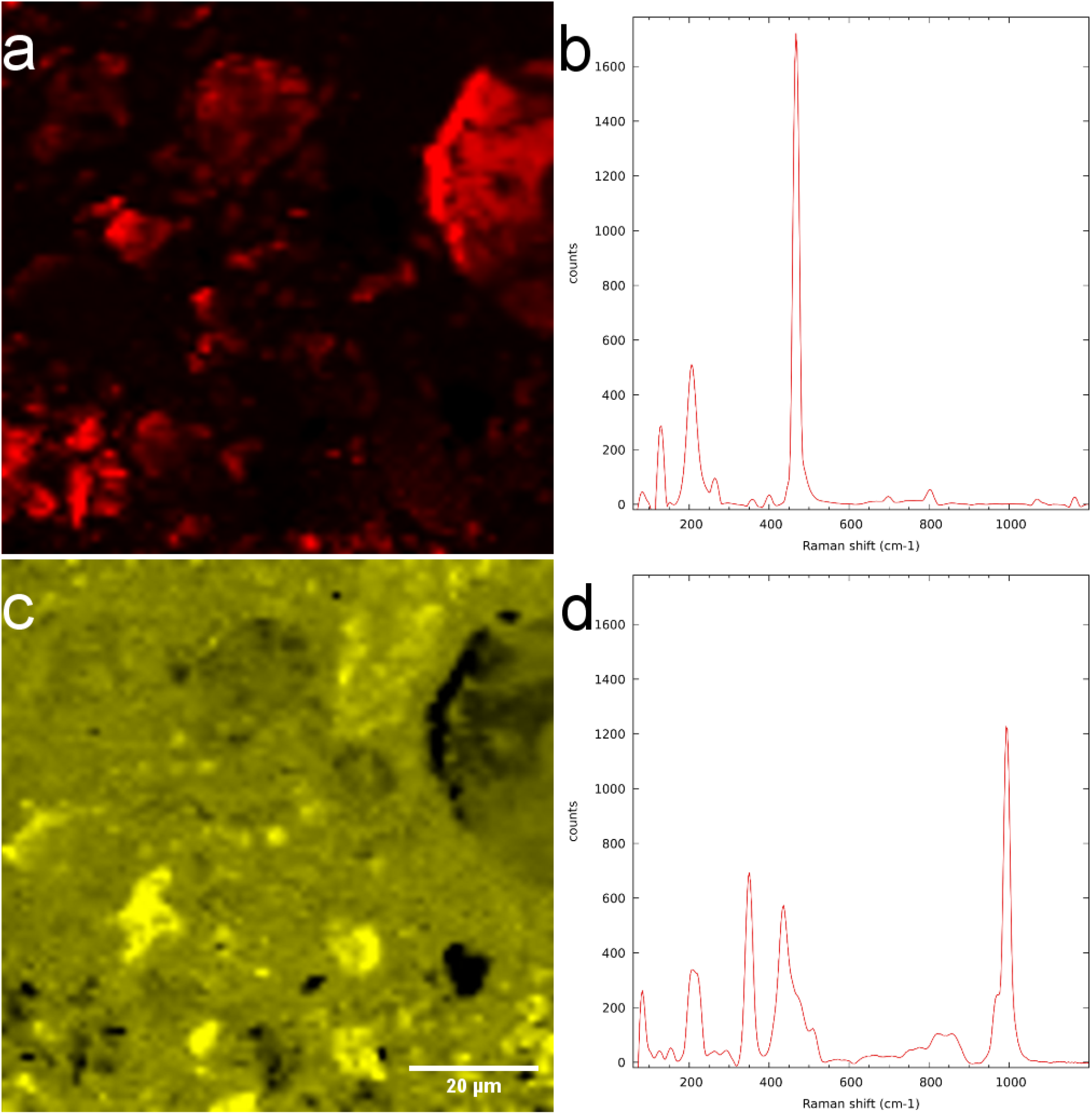
µ-Raman measurement of LR-white resin-embedded soil. a) Quartz-distribution map and b) spectrum. c) Mixture of zircon and aluminium-containing minerals and their d) spectra.

#### The spatial distribution of bacteria and CARD-FISH

On the LR white embedded sample, we could successfully identify bacterial aggregates within the soil using CARD-FISH (Figure 8). Bacteria fluoresced in DsRed channel is false colored in red orange under excitation wavelength of 590 nm after successful hybridization, making it easy to distinguish bacteria from the native auto-fluorescence of the soil. (See SI for more Figures). In contrast, the DAPI staining was non-specifically binding to soil components and resin, making difficult cell identification. However, for each positively hybridized microbial cell, the DAPI signal was verified to coincide with the Alexa594 signal. Bacteria were often observed as clusters or agglomerations rarely in the vicinity of root sections and mostly in the vicinity of quartz minerals (Fig 8). As stated in literature,^15, 17^ it is evident that nuclear stains are not capable of identifying bacteria in soil and specialized CARD-FISH methods are required for a precise identification. However, single microbial cells are hard to visualize due to high background fluorescence, particularly of the roots and associated minerals, we therefore cannot exclude the presence of these in the root surroundings. Correlative EDX analysis (Figure 8d) shows the chemical composition of the minerals in the vicinity of the microbial hot spots. However, further investigations are necessary to determine the type of minerals which the bacteria colonize.

**Figure 8:**
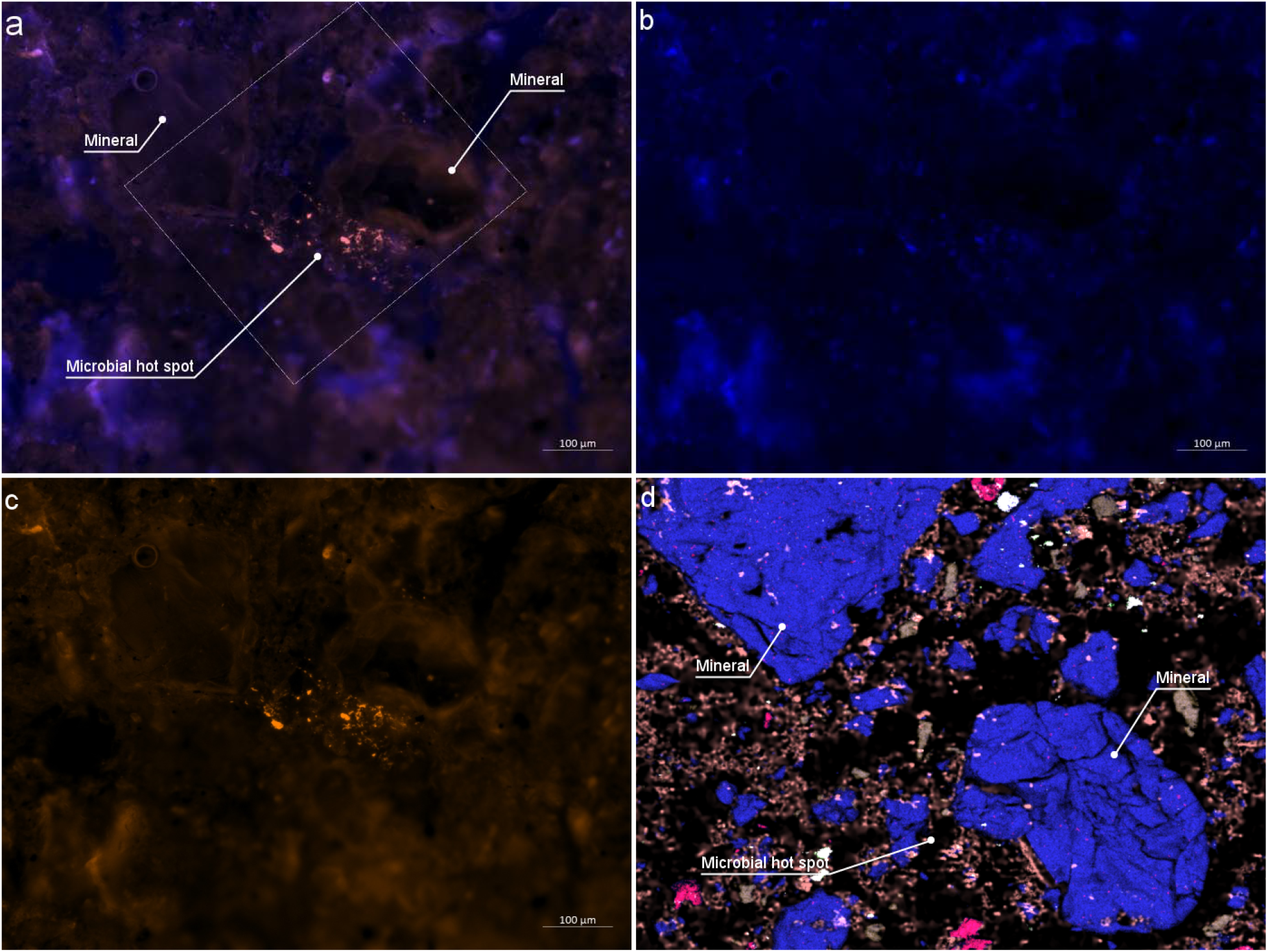
CARD-FISH stained bacteria colonizing on Rhizosphere mineral. a) Epifluorescence micrograph of combined DAPI and DsRed channel. Outlined square shows the area investigated by EDX analysis b) DAPI channel c) DsRed channel d) Same area imaged by EDX. Two larger minerals indicate Si rich minerals where bacterial aggregates are identified

**Figure 9:**
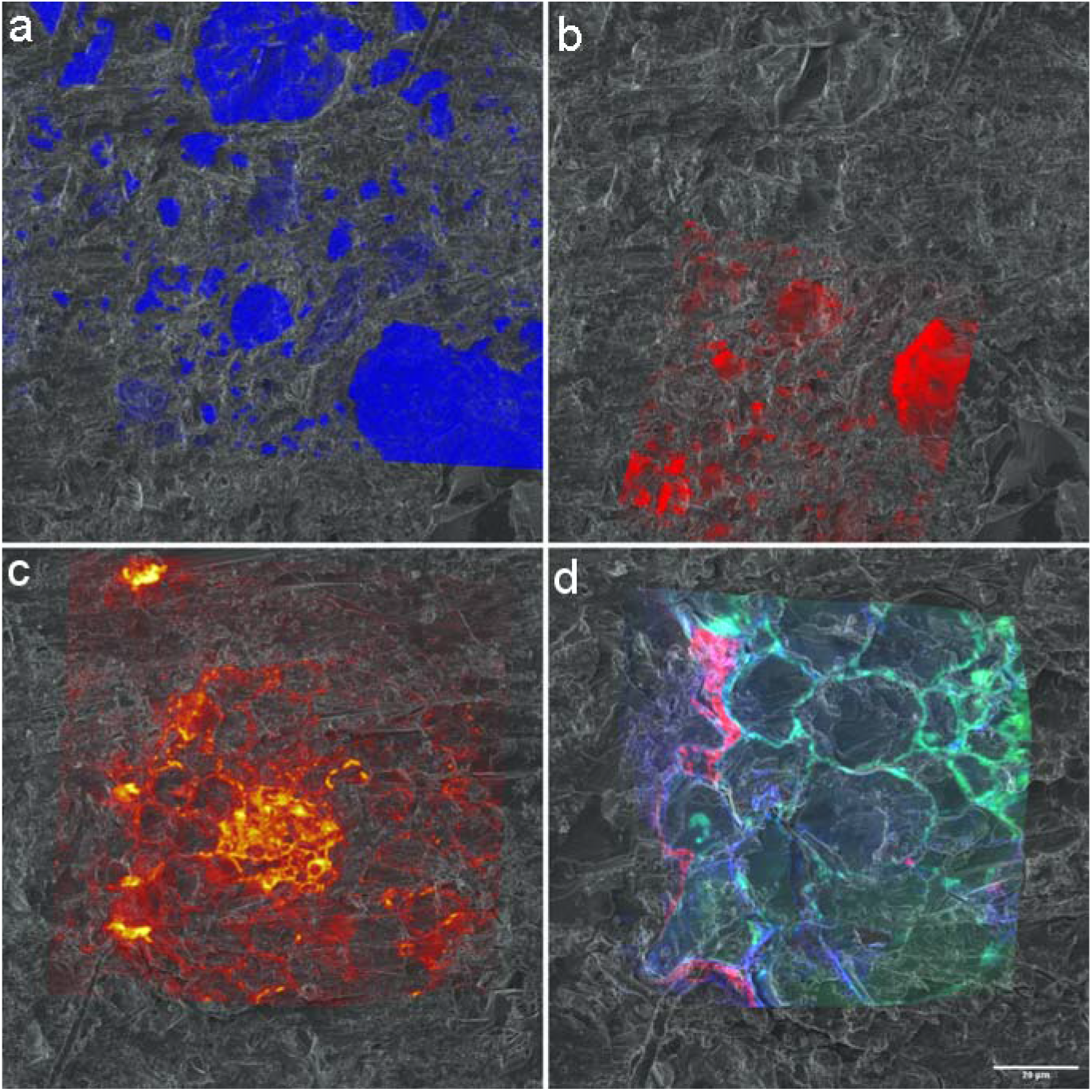
Chemical Micrographs registered to HIM Micrographs a) EDX spectra of Si registered onto HIM b) Raman spectra resisted on to HIM c) P spectra of ToF-SIMS registered onto HIM. d) ^12^C^14^N^-, 31^P^-^ and ^32^S^-^ distribution maps from nanoSIMS registered onto HIM.

The success of the CARD-FISH experiment strongly depends on the temperature at which the resin is cured. Therefore, we cured the resin at 48 °C, and at 60 °C to test the hybridization success. CARD-FISH were only successful at 48 °C, which is well below the standard polymerization temperature of 60 °C. Polymerization of LR white is an exothermic reaction and curing at 60 °C can further increase the embedding temperature leading to potential degradation of the bacterial ribosomes, also have a negative effect of probe permeability through the strongly polymerized resin. These results are in-line with our observations, where CARD-FISH experiments were not successful with the samples cured at 60 °C, instead hybridization was observed only at 48 °C. Thus, if correlation with CARD-FISH is desired, temperature of 48 °C should be used for curing and found to be compatible for high vacuum microscopy.

Different microscopy techniques may alter the surface topography due to beam-sample interactions and lead to artefacts. (See SI) As such, it is important to setup the characterization cascade starting with techniques that cause minimum damages while measurement. This will allow to acquire the maximum set of correlative data with minimum artefacts. Here, we suggest a characterization cascade starting with light microscopy for mapping, followed by CARD-FISH and epifluorescence microscopy. Then the following the order HIM → ToF-SIMS → SEM/EDX → nanoSIMS → µ-Raman for high-resolution characterization. HIM and ToF-SIMS being surface sensitive techniques, to be completed at earlier stages. Starting from SEM sample made conductive by depositing a conductive metal layer, leading to minor surface modifications. NanoSIMS is a destructive technique providing hard ionization of sample material up to single atomic ions for analysis of the isotopic composition resulting major pits in the sample. Hence it is to be carried out after SEM/EDX. Raman scanning is the last as it is time consuming and the laser can cause the resin evaporation, but minerals are unlikely to be altered by other techniques.

### 3.3 Image Registration with the Correlia plugin in Fiji

For a mechanistic understanding of complex patterns of interactions between soil, microbiome and the roots using a correlative approach, multiple images acquired by various techniques needs to be successfully registered and to be analyzed. For the image registration we have used helium ion micrographs as the base image to register all high-resolution chemical micrographs as the first step as similar features can be recognized. Then HIM was registered on to the optical micrograph based on common surface features. (See SI) HIM is an excellent alternative to optical or SEM micrographs as it can be used as early characterization tool in this workflow providing excellent resolution and quick image acquisition compared to other techniques. Figure 10 demonstrates the registration of a) SEM-EDX, b) µ-Raman, c) ToF-SIMS, and d) nanoSIMS micrographs onto HIM micrograph. Our results demonstrate HIM is a well-suited for non-quantitative imaging of the embedded rhizosphere sample to prepare treasure maps of the interested RoI and image registration. However, in a correlative workflow involving X-Ray micro-µCT, BSE is a promising candidate for 2D-3D image registrations due to their similar contrast.

## Conclusions

Here we presented a new embedding method that allow comprehensive correlative microscopic approach which make way for a synergistic understanding of complex soil-plant-microbe interactions via multiple microscopic and micro-analytical techniques. Introduced LR white embedding and waterjet cutting showed capability of preparing the rhizosphere soil matrix compatible for high-resolution chemical microscopy and identification of bacteria by CARD-FISH, which were not combined before in a correlative approach. Now vital research studies can be carried out using a variety of instruments with specialist analytical capabilities in combination, covering all components of rhizosphere: soil-root-microbes. Overall, this method allows to comprehensively study the spatio-temporal organization of nutrients and microbes in the rhizosphere at µm to nm scale and we hope this will make a platform for mechanistic understanding of complex patterns of interactions between roots, microbiome and the soil in a correlative microscopy approach in the future.

## Author contributions

CB-Developed the embedding method, HIM imaging, epifluorescence imaging, ToF-SIMS analysis, image registration, data analysis and wrote the initial draft

YD-Analyzed the Raman data and contributed to manuscript editing

MS-HIM imaging, Raman and SEM/EDX analysis, image registration and contributed to manuscript writing

HS-ToF-SIMS and nanoSIMS experiments and data analysis, contributed to manuscript writing

HR-Acquired funding, managed the overall project, contributed to manuscript editing and project supervision

NM-CARD-FISH experiments, project supervision and contributed to manuscript writing

All authors read and agreed on the final version of the manuscript.

## Funding

*This work was conducted within the framework of the priority program 2089, “Rhizosphere spatiotemporal organization-a key to rhizosphere functions” funded by the Deutsche Forschungsgemeinschaft (DFG, German Research Foundation) – Project number RI-903/7-1. Research work of Yalda Devoudpour was supported by Deutsche Forschungsgemeinschaft Integration of Refugee Scientists and Academics. Open access was supported by UFZ library*.

## Acknowledgement

We acknowledge the analytical facilities of the Centre for Chemical Microscopy (ProVIS) at the Helmholtz Centre for Environmental Research, Leipzig, Germany, which is supported by the European Regional Development Funds (EFRE - Europe funds Saxony) and the Helmholtz Association. The authors are grateful to Frank Hochholdinger for the supply of *Zea mays* seeds. Authors thank Eva Lippold, Dr. Steffen Schulter and Prof. Doris Vetterlein at the Helmholtz Centre for Environmental Research, Halle for conducting and coordinating soil column experiments. Authors acknowledge the technical contributions from Katja Nerlich, Jasmin Voight for CARD-FISH experiments and Dr. Hugo Berthelot for contributions during sampling. The development of LANS software by Dr. Lubos Polerecky (Utrecht University) and his kind application support are greatly acknowledged. Finally, authors thank for the Peter Portius, Daniel Karmowski and Manuel Kositzke at UFZ workshop for water-jet cutting.

## Notes

### Competing Interest Statement

The authors have declared no competing interest.

### Summary of Updates

Typographical errors in the abstract and method section was corrected.

