## Supporting Information for "High-Resolution Chemical Mapping and Microbial Identification of Rhizosphere using Correlative Microscopy"

Running Title: Comprehensive correlative microscopy approach for rhizosphere characterization

Chaturanga D. Bandara*, Matthias Schmidt, Yalda Davoudpour, Hryhoriy Stryhanyuk*, Hans H. Richnow, Niculina Musat

Department of Isotope Biogeochemistry, Helmholtz-Center for Environmental Research (UFZ), Permoserstraße15, 04318 Leipzig, Germany

### Application of multiple micro analytical tools in soil research

***HIM*** is an extremely surface-sensitive (probing only a few nm below the surface), high-resolution (down to 0.25nm) microscopy technique with numerous applications in the life-sciences. [1-3] The technique allows for investigating insulating samples owing to efficient charge-compensation with an electron flood-gun, [1, 4] On top of that, HIM imaging is operating at beam-currents in the sub-pico-amp range which in turn means that only a few tens of ions are being implanted per pixel during image acquisition causing only a minimum damage to the sample.[2-3, 5] To the best of our knowledge, to-date only one publication in soil research exists where HIM was used to study mineral-organic interaction. [3] Therefore, adding HIM to the toolbox of correlative analysis of the rhizosphere with high-resolution 2D imaging techniques provides additional insight to soil biochemical processes at nanoscale of intact rhizosphere.

***SEM*** is a well-established surface-analytical technique in soil-science and geology. Whilst the commonly used secondary electron (SE) imaging provides high lateral resolution and surface sensitivity no chemical information about the imaged material can be gained. In contrast, a material contrast increasing with an electron density is obtained when using electron acceleration energies of >=10 kV and detecting back-scattered electrons (BSE).[5] In resin-embedded soil this allows for separating resin and organic matter from mineral particles. Furthermore, element-specific chemical information of the soil sample, in particular to identify mineral particles, can be obtained when the SEM is combined with an EDX.[6]

***ToF-SIMS*** allows for studying the molecular composition of heterogeneous organic/inorganic samples with about 100 nm spatial resolution and simultaneous detection of molecular fragments with mass-resolving power >5000 (MRP=dM/M) in a broad mass range up to ~10 kDa.[7-8] The application of cluster-ion sources in a ToF-SIMS experiment reduces molecular fragmentation and allows for high-resolution 3D analysis of organics without having to destroy the spatial organization of the sample by chemical extraction. This renders ToF-SIMS a potential technique to study biomarkers in complex mineral-organic soil specimens.[9-12] Recent trend in advances of complex ToF-SIMS data analysis including Principal component analysis (PCA), multivariate curve resolution-altering least squares (MCR-ALS), G-SIMS[13] and artificial intelligence methods like neural networks and novel informatics-based methods[11, 14-20] will eventually lead to identify various molecules in a complex mixture of organics in rhizosphere sample in the future.

***nanoSIMS*** allows for elemental/isotope-resolved mapping of sample surface with simultaneous detection of ion counts for up to 7 isotopes (nanoSIMS 50L), lateral resolution down to 50 nm and mass-resolving power (MRP=M/dM) above 8000.[21] Combining nanoSIMS with Fluorescence and or Halogen *In-situ* Hybridization (FISH/HISH) and Stable Isotope Probing (SIP) allows for phenotypic identification of bacteria and quantitation of their metabolic activity in complex environmental samples.[22-24] The recently commercialized RF-plasma source of $O^-$ ions makes nanoSIMS feasible for high-resolution mapping of trace elements in biological systems.[25]

**µ-Raman** is based on the inelastic scattering of the monochromatic light at the surface of a sample and allows for spatially resolved molecular identification based on molecules unique vibrational characteristics.[26] µ-Raman analysis does not require elaborative sample preparation steps which makes it easily applicable on a variety of samples from geological specimens to the characterization of soil, soil organic matter and carbon flows.[27-29] Recent technical improvements extended its application in earth- and life-sciences, particularly for microbial cells analysis and isotopes.[30-31] Thus, µ-Raman contributed to the identification and classification of soil bacteria and their metabolites in combination with stable isotope labelling and other single cell techniques such as namoSIMS[32-33] e.g. spotting nitrogen fixing bacteria in soil community using $^{15}N$ stable isotope labelling.[30, 34]

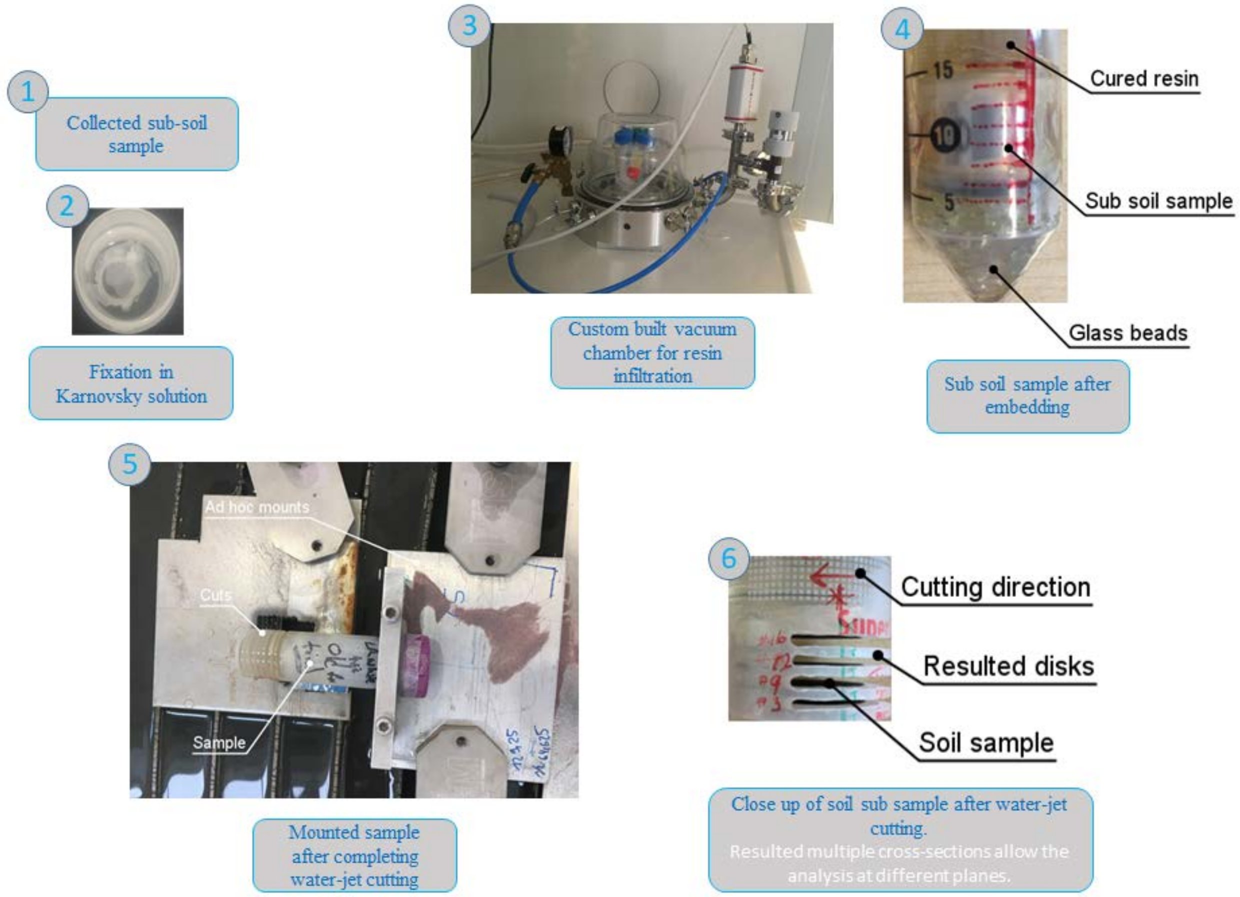


*Supporting Figure S 1: Pictures from individual steps of the embedding method*

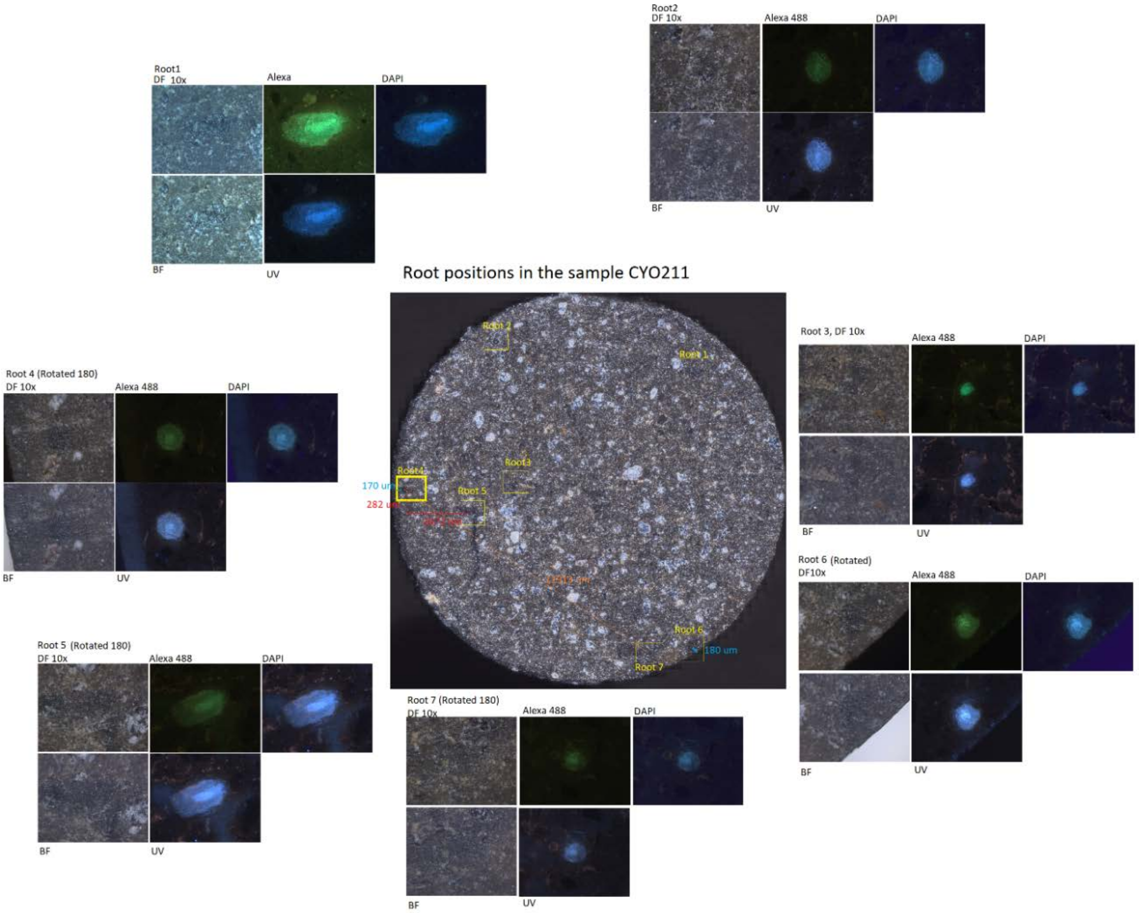


*Supporting Figure S 2: Identification of RoI with Epifluorescence microscopy and prepared treasure map for high resolution microscopy*

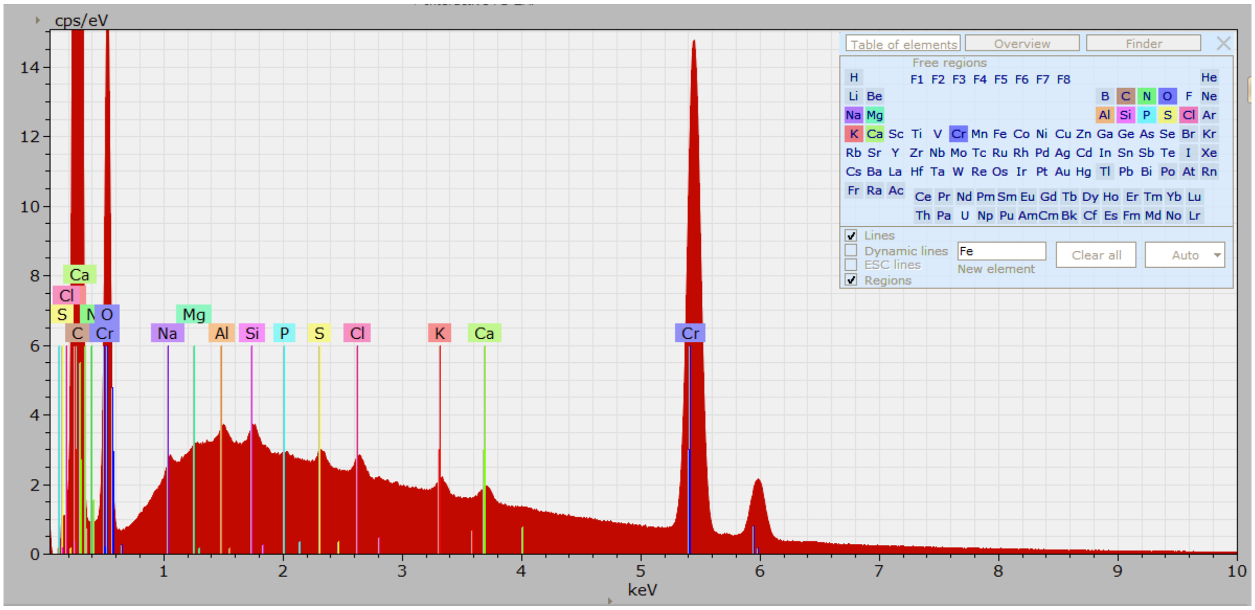


*Supporting Figure S 3: EDX Spectra of LR white*

*Supporting Table 1: Water-jet cutting parameters operated at 3700 bar pressure*

| Setting Number | Feed Speed in mm/min | Particle consumption rate g/min |
|---|---|---|
| 1 | 54.7 | 400 |
| 2 | 54.7 | 400 |
| 3 | 27.35 | 400 |
| 4 | 5.55 | 400 |
| 5 | 125.9 | 350 |
| 6 | 89.2 | 350 |
| 7 | 65.3 | 500 |
| 8 | 32.6 | 500 |
| 9 | 6.63 | 500 |
| 10 | 220.8 | 350 |
| 11 | 111.4 | 350 |
| 12 | 25.8 | 400 |
| 13 | 12.9 | 400 |
| 14 | 16.9 | 400 |
| 15 | 14.7 | 450 |
| 16 | 7.3 | 450 |
| 17 | 169.7 | 350 |
| 18 | 84.8 | 350 |

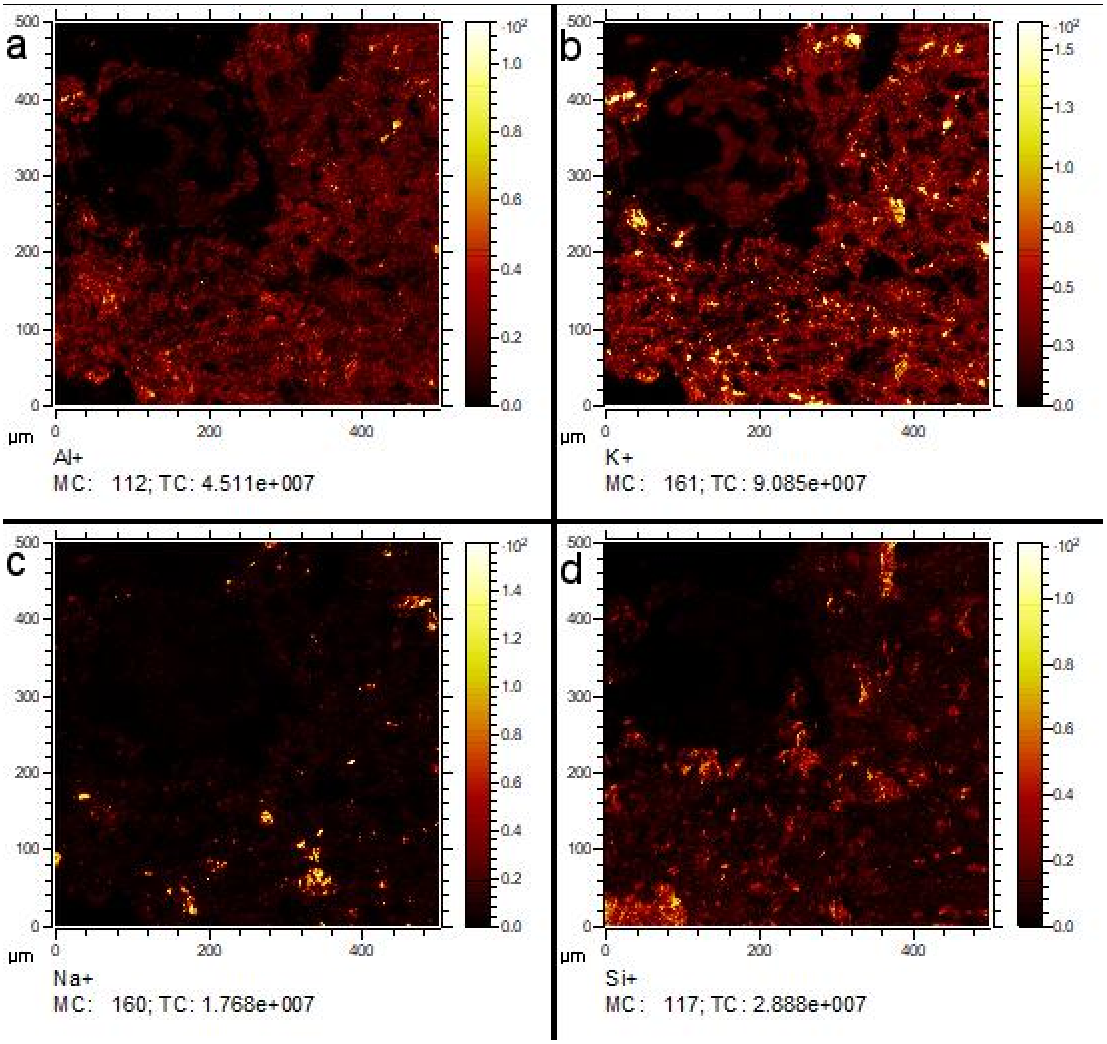


*Supporting Figure S 4: ToF-SIMS of the rhizosphere in positive extraction mode. Al, K, Na and Si minerals are distinguished by Multivariate analysis. Root area shows no signal and hence dark.*

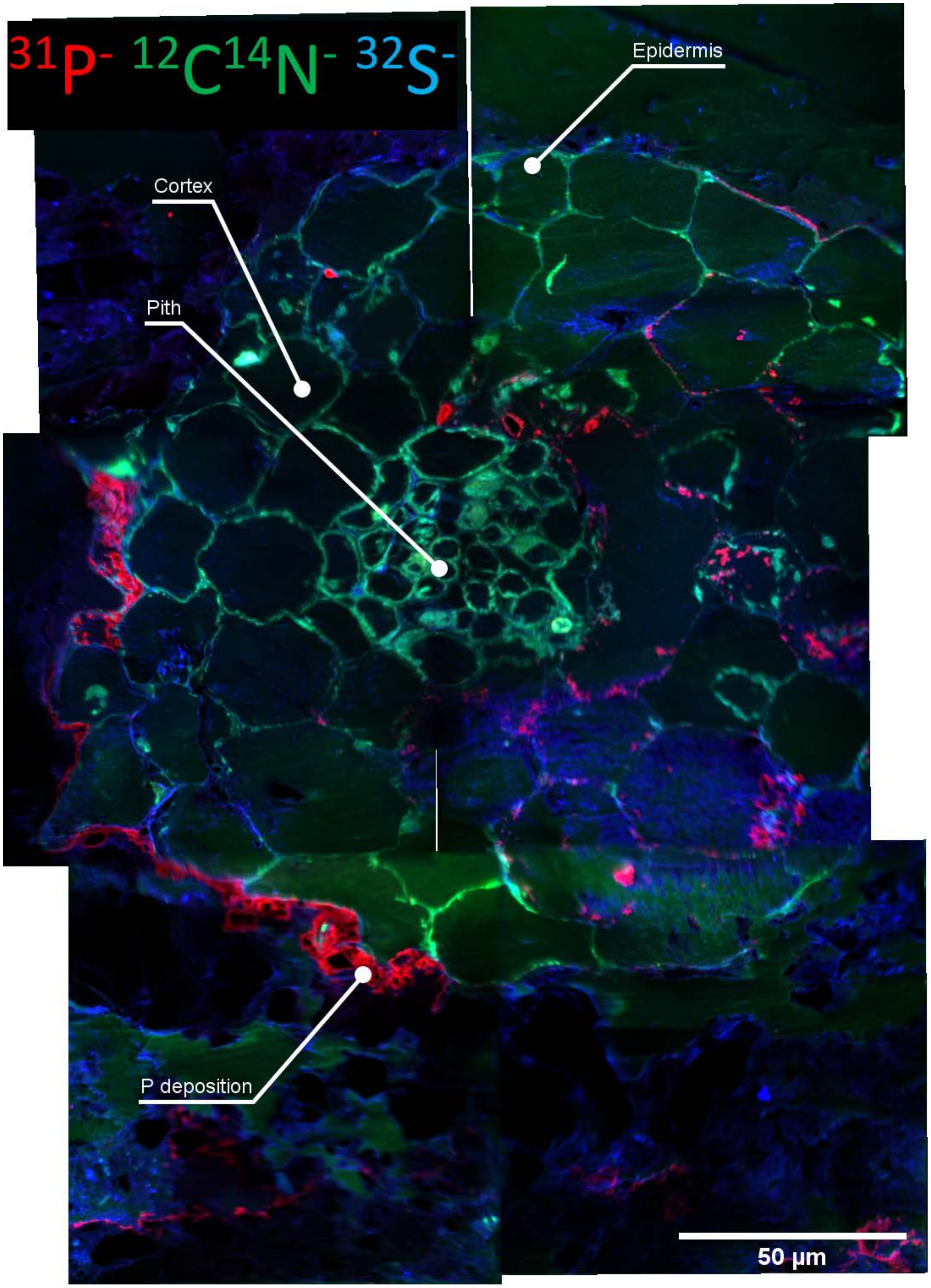


*Supporting Figure S 5: nanoSIMS mosaic of rhizosphere showing $^{31}P^{-}$, $^{12}C^{14}N^{-}$ and $^{32}S^{-}$ distribution. Image is reconstructed by combining 7 individual FoVs. Individual image is 95 µm FOV. P deposition is prominent on the epidermis of bottom left quarter of the root. In the adjacent cortex P is not detected. When P is not detected on epidermis, it is detected on the adjacent cortex cells.*

*Supporting Table 2: Characteristic  Raman peaks on Zircon and aluminium ammonium sulfate*

| Sample | Band position (cm-1) | Suggested assignment |
|---|---|---|
| Zircon[35] | 1000 | Si-O stretching |
| | 436 | Si-O bending |
| | 350 | External mode |
| Aluminium ammonium slufate[36] | 995 | $SO_4$ |
| | 600 | $SO_4$, $AlO_6$, Al-OH |
| | 450 | Lattice modes of ($AlO_6$), (Al-$H_2O$), ($SO_4$), ($NH_4$) |

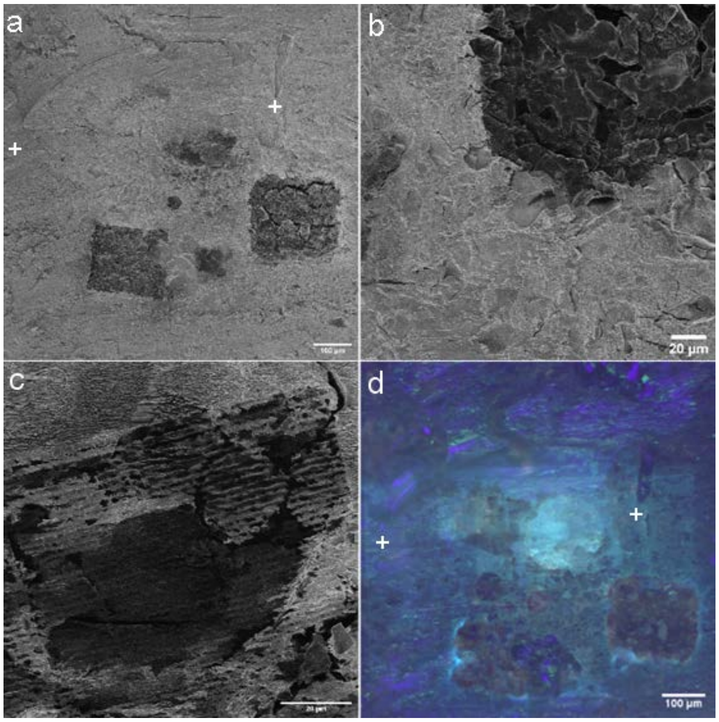


*Supporting Figure S6: Sample damage after correlative microscopy. a-c) HIM micrographs show the overview of the sample after multiple imaging. b) Damaged caused by 30mW power laser with 425 nm wavelength. c) Damaged caused by 10 mW power laser with 785 nm wavelength. Damage is much less compared to b. d) Epi-fluorescence micrograph show the difference in fluorescence in the sample after characterization by multiple microanalytical tools. Areas scanned with Raman microscope are not fluorescent.*

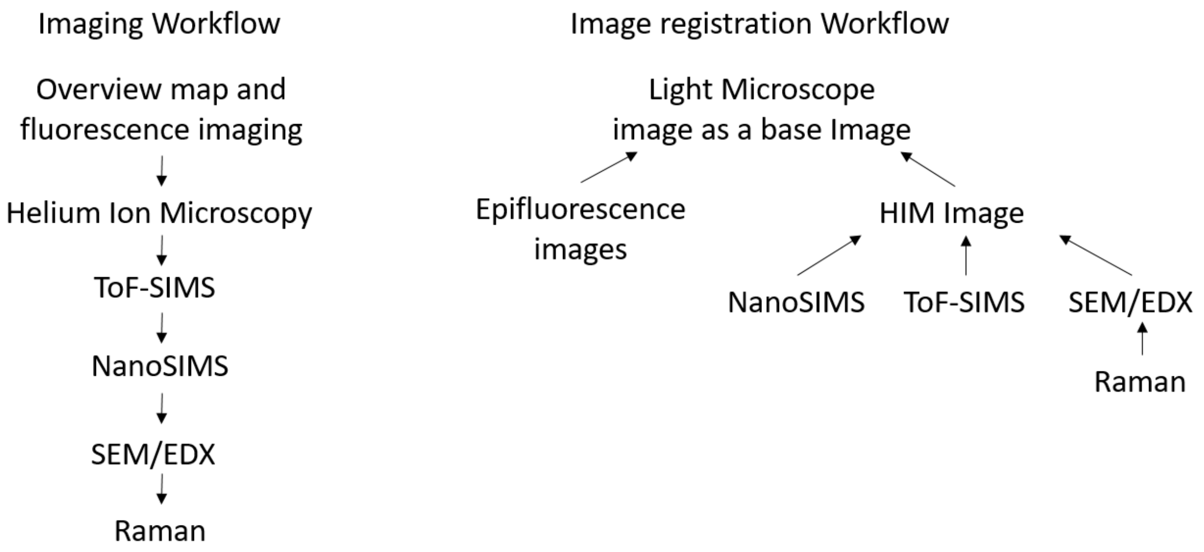


*Supporting Figure S 7: Image acquisition and Image registration workflows. Imaging workflow shows the recommended image acquisition. Image registration suggests the high resolution images are to be first registered on to HIM, Raman images to the EDX map from SEM and then HIM image to be registered on to light micrographs.*

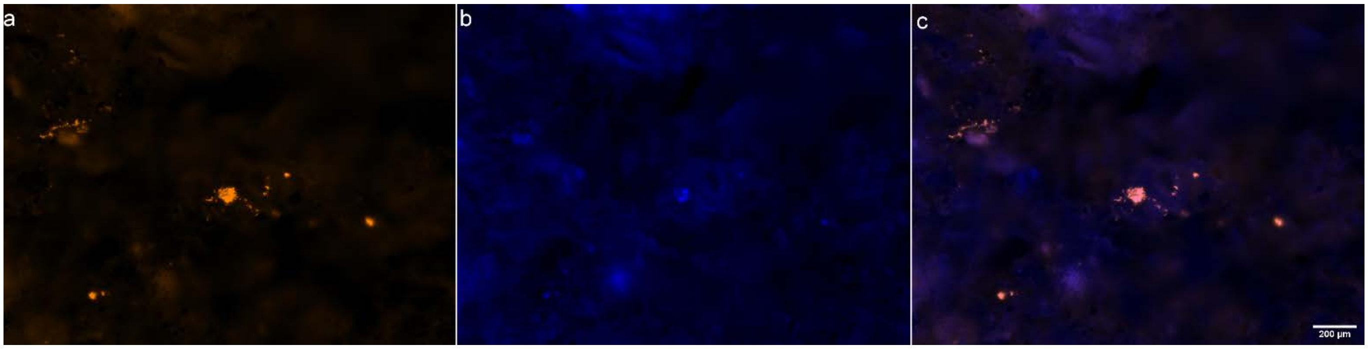


*Supporting Figure S 8: CARD-FISH showing multiple bacterial colonies of embedded rhizosphere a) Ds Red b)DAPI c) Two channels combined*

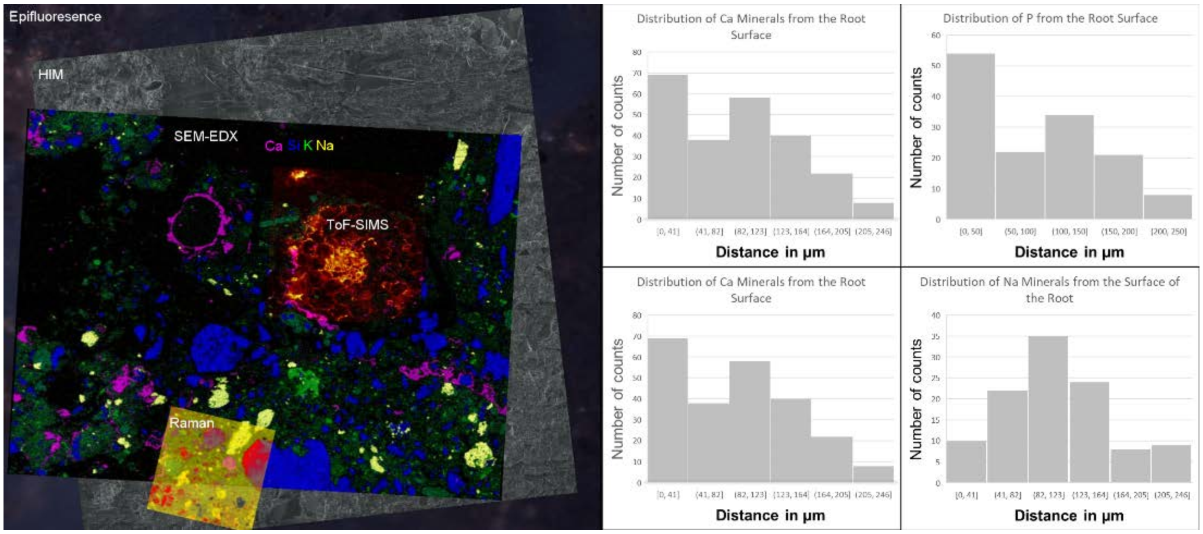


*Supporting Figure S 9: Distribution of elements in the rhizosphere. Distance is calculated from the surface of the root.*

Here, we demonstrate a possible analysis of registered EDX and TOF-SIMS data using the Correlia Plugin, showing the elemental distribution on a 2D plane from the epidermis of the root. As EDX cannot distinguish root cell wall due to the similar composition as of resin, use of ToF-SIMS could overcome the analysis difficulties, which is a highlight of correlation of multiple imaging techniques. This is only a plausible demonstration on various data analyses could be done with an availability of a method which can characterize the all-important components of the rhizosphere.
